# Mediodorsal thalamic nucleus mediates resistance to ethanol through Cav3.1 T-type Ca^2+^ regulation of neural activity

**DOI:** 10.1101/2023.09.25.558585

**Authors:** Charles-Francois V. Latchoumane, Joon-Hyuk Lee, Seong-Wook Kim, Jinhyun Kim, Hee-Sup Shin

## Abstract

Thalamocortical activity is known to orchestrate sensory gating and consciousness switching. The precise thalamic regions involved, or the firing patterns related to the unconsciousness remain unclear. Interestingly, the highly-expressed thalamic T-type calcium currents have been considered as a candidate for the ionic mechanism for the generation of thalamo-cortically-driven change in conscious state. Here, we tested the hypothesis that Ca^v^3.1 T-type channels in the mediodorsal thalamic nucleus (MD) might control neuronal firing during unconsciousness using Ca^v^3.1 T-type channel knock-out (KO) and knock-down (KD) mice under natural sleep and ethanol-induced unconsciousness. During natural sleep, the MD neurons in KO mice showed general characteristics of sustained firing across sleep stages. We found that KO and MD-specific KD mice showed enhanced resistance to ethanol. During ethanol-induced unconscious state, wild-type (WT) MD neurons showed a significant reduction in neuronal firing from baseline with increased burst firing, whereas Ca^v^3.1 KO neurons showed well sustained neural firing, within the level of wakefulness, and no burst firing. Further, 20 Hz optogenetic and electrical activation of MD neurons mimicked the ethanol resistance behavior in WT mice. These results suggest that maintaining MD neural firing at a wakeful level is sufficient to induce resistance to ethanol-induced hypnosis in WT mice. This work has important implications for the design of treatments for consciousness disorders using thalamic stimulation of deeper nuclei including the targeting of the mediodorsal thalamic nucleus.

**GRAPHICAL ABSTRACT:** 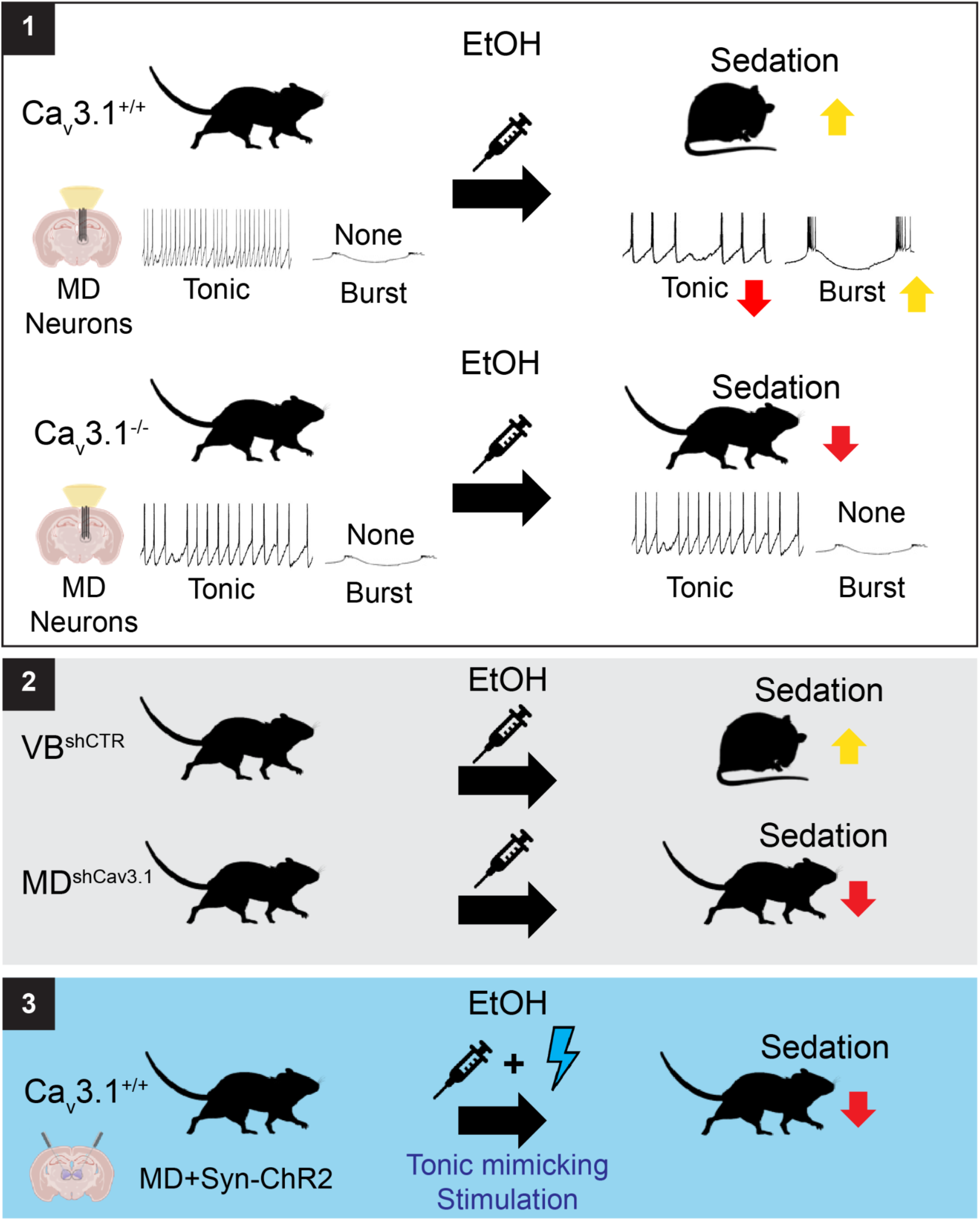

## INTRODUCTION

Drug-induced unconsciousness can be achieved using numerous types of anesthetics with varying modes of action ^1,2^. Ethanol, one of the most frequently abused drugs in human society, can induce sleep-like loss of consciousness at high doses ^3^. While possible neuropharmacological and neural correlates of ethanol sedation have been proposed using *in vitro* and *in vivo* methods ^4–7^, recent studies have highlighted the slowing of thalamocortical-driven rhythms as a potent marker of unconsciousness ^8,9^. However, the region and the mechanism linked to the thalamic modulation during ethanol induced unconsciousness remains poorly understood.

Physiological correlates of thalamocortical rhythmic activities and consciousness state of the brain have been studied extensively^10–13^. T-type calcium channels are known generators of thalamocortical rhythms, through the modulation of cell excitability and rebound burst firing^14,15^. During sleep, the transition from wakefulness to unconsciousness is associated with membrane hyperpolarization of thalamic neurons ^12,16^. Similarly, It has been shown that in ethanol sedation, as in natural sleep or absence seizure, the loss of consciousness is characterized by a switch from tonic to burst firing in thalamic neurons, which involves GABAergic inhibition-driven de-inactivation of Ca^v^3.1 T-type channels resulting in slow oscillatory response of the thalamocortical network ^7,17–19^. Acute intoxication at high doses of ethanol ^20,21^ induces both slow oscillations in the delta-theta frequency range and a loss of righting reflex (LORR), a classical proxy to assess the loss of consciousness. It has been shown that mice lacking global or thalamic Ca^v^3.1 showed altered slow oscillations and sleep architecture ^22,23^; delayed sleep induction under several anesthetics (i.e. isoflurane, halothane, sevoflurane and pentobarbital) ^24^; and increased resistance to drug-induced absence seizure ^13^. Notably, the absence or blockade of Ca^v^3.1 resulted in an increased preference for ethanol consumption and novelty-seeking behavior ^20,21^. In the current studies we investigate the role in brain state modulation of Ca^v^3.1-mediated T-currents during ethanol-induced sleep.

The thalamus is one of the major regions expressing Ca^v^3.1 T-type calcium channels ^25^ and holds a central role in information-transmission and integration ^26^. *In vitro* and *in vivo* studies using genetically modified mice have revealed that Ca^v^3.1 T-type channels play a key role in the genesis of thalamocortical rhythms, such as 3Hz spike-and-wave discharges (SWD), a signature of absence seizures ^13,27^ and delta waves ^16,28,29^. Previous investigations on thalamic control of consciousness revealed that nuclei within the dorsal medial thalamus (dMT) hold an important modulatory function in the interaction of attention and arousal ^9,30^. Particularly, centromedian (CM) thalamic nucleus, and not ventrobasal nucleus (VB), showed rapid shifts in local field potential (LFP) preceding brain state transitions such as NREM and propofol-induced anesthesia ^8^. The paraventricular thalamic nucleus (PVN) showed critical involvement in wake/sleep cycle regulation ^33^. The centrolateral (CL) thalamic nucleus has been implicated in the modulation of arousal, behavior arrest ^31^, and improvement of level of consciousness during seizures ^32^. Notably, the direct electrical stimulation of the intralaminar nuclei (ILN) and, in particular CL, promoted hallmarks of arousal and awakening in primate under propofol and ketamine propofol anesthesia. The MD, a sub nucleus of dMT, on the other hand, has only recently been implicated in disorders of consciousness ^33^ and ketamine/ethanol-induced loss of consciousness^34^ through the alteration of thalamo-cortical functional connectivity. In anesthetized primate, the stimulation of ILN and MD increased arousal and wakefulness score^35^. However, several key questions still remain to be answered: 1) Is there a specific role for MD Ca^v^3.1 T-type calcium channels in the control of ethanol-induced loss of consciousness? 2) Whether Ca^v^3.1 T-type calcium channel driven neuronal firing pattern has any role in the control of consciousness?

In this study, we identified that KO and MD specific silencing of Ca^v^3.1 T-type calcium channels result in increased ethanol resistance in mice. Using single unit recordings, we compared MD activity of WT and KO mice while the mice transitioned from conscious to unconscious state and found that the KO mice showed more sustained MD activity whereas the WT mice showed clearly reduced MD activity. Furthermore and consistently with their resistant phenotype, KO mice showed sustained MD firing, well within the wakefulness level, under ethanol consumption. Finally, we demonstrate that both the optogenetic and electrical stimulations in MD, mimicking the sustained firing pattern of knock-out mice, were sufficient to induce the increased ethanol resistance in WT mice. These results reveal a causal control of brain state by MD during ethanol-induced unconsciousness and with an underlying neural mechanism governed by Cav3.1 T-type calcium channels.

## RESULTS

To understand the role of T-type Ca2+ channels in modulating the consciousness level, we compared the ethanol resistance between WT and Ca^v^3.1^−/−^ KO littermates. We used the forced walking task (FWT; Fig. 1-A), an analog to the LORR assay, which enables a continuous and high-temporal resolution assessment of the loss of movement ^36^ (LOM). Moreover, the FWT objectively measures the latency to and duration of the first LOM, but also the total time spent in LOM using automatized analysis of video confirmed by electromyograms (EMGs) or accelerometer recordings (Fig. 1-B and S1; see Methods). The continuously running treadmill (6 cm/s) ensures a normalized behavior within and between animals before injection (i.e., baseline forced walking) and allows for reduced intervention from experimenters for the monitoring of both electrophysiological (Fig. S1, upper panel) and analyzed behavioral (Fig. S1, lower panel).

**Fig. 1:**
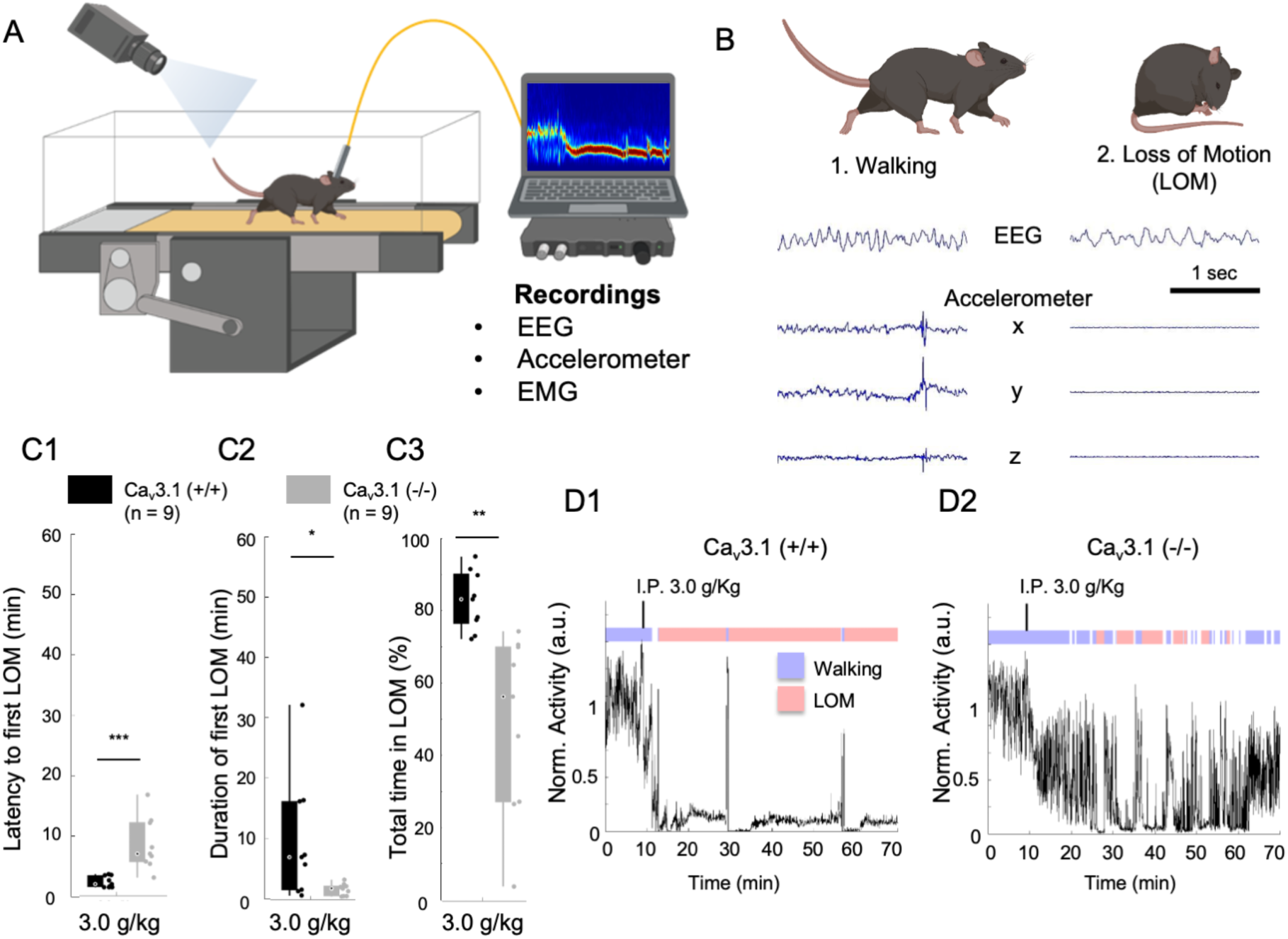
Mice lacking Ca^v^3.1 showed increased ethanol resistance on the forced walking task. (A) The schematic of the FWT setup. Mice are habituated and trained on a constantly moving treadmill (6 cm/sec). Following a baseline walking recording (∼10 min), the mouse is carefully picked up and injected with ethanol (i.p.). Once placed back on the treadmill, the loss of consciousness is evaluated using normalized moving index using either video analysis (differential pixel motion), on-head accelerometer-based motion, or neck electromyograms over a period of 60 min. (B) Representative EEG (parietal) and 3-axis accelerometer (Acc) traces for walking and LOM in a Ca^v^3.1 (+/+) mouse. (C) Quantification for the latency to first LOM (fLOM; i.e. delay between I.P. Injection and loss of motion; C1), the duration of the fLOM (C2) and the total time spent in LOM state (C3) over a recording duration of 60 min following 3.0 g/Kg I.P injection of ethanol in Ca^v^3.1 (+/+) and Ca^v^3.1 (-/-) mice; data is represented as boxplot wish individual mice as scatter plot. * is for p<0.05, ** is for p<0.01 and *** is for p<0.001. (D) Representative normalized motor activity over time for Ca^v^3.1(+/+) (D1) and Ca^v^3.1 (-/-) (D2) mice post I.P. injection of 3.0 g/Kg. Blue and red boxes above the graph indicate the state interpretation for walking and LOM, respectively.

### The lacking Ca^v^3.1 increases ethanol resistance in mice

Mice lacking Ca^v^3.1 exhibited delayed anesthetic induction ^13,24^ and impairment in maintenance of low conscious level ^22,23^ as well as increased ethanol preference ^21,34^. We tested the sensitivity of Ca^v^3.1 null mutant mice for various acute hypnotic doses of ethanol.

We confirmed that the lack of Ca^v^3.1 resulted in a more delayed and fragmented LOM (Fig. 1-C1 and C2), and a reduction in the total time spent in LOM compared to Ca^v^3.1 WT mice (Fig. 1-C3). We observed that Ca^v^3.1 null mutant mice showed increased latency to and decreased duration of the first episode of loss of motion (fLOM) for ethanol injection doses of 2.0, 3.0 and 4.0 g/Kg (Fig. S2). The total time spent in LOM during one hour of recording was also significantly reduced (Fig. S2-A3 and -B3) compared with WT mice. Two-way analysis of variance (ANOVA) showed a significant effect for the main factor genotype and dose for the latency to and duration of fLOM, and total time in LOM (Table S1), indicating a dose dependency in both wild and mutant mice. In particular, an I.P. Injection of 3.0 g/Kg induced a significant difference between wild and mutant mice in latency to (t(16) =-4.1965, p = 0.002; Student t-test) and duration of fLOM (t(16) =2.3908, p = 0.0294; Student t-test), and total time spent in LOM (t(16) =3.9065, p = 0.0012; Student t-test).

These results indicate that ethanol induces a dose-dependent sedative effect on mice and Ca^v^3.1 mutant mice had an increased resistance to ethanol sedation compared to WT mice.

### Ca^v^3.1 silencing in the MD, but not VB, increased ethanol resistance in mice

A great majority of Ca^v^3.1 expression is found in the thalamic region ^25^ and was shown to specifically correlate with the modulation of thalamocortical-related rhythms and stability of sleep level ^22,23^. To identify a possible role of the thalamic region in ethanol resistance, we knocked down the expression of Ca^v^3.1 in the MD and a ventral basal nucleus (VB) region, two regions possibly involved in a thalamic control of consciousness ^33,34,37^, using a lentivirus (LV)-mediated short hairpin (shRNA) delivery.

We found that compared to shControl injected mice, shCa^v^3.1 knock-down of MD resulted in an increased latency to (Fig. 2-A1; t(27) = -3.0045, p = 0.0057; Student t-test) and duration of (Fig. 2-A2; t(27) =2.1448, p = 0.0411; Student t-test) fLOM, and total time spent in LOM (Fig. 2-A3; t(27) =2.6641, p = 0.0128, two-tailed test) for 3.0g/Kg I.P. Injection of ethanol. However, we found that compared to shControl-injected mice, Ca^v^3.1 KD of VB did not change the latency to (t(8) =-1.0093 p = 0.3423, two-tailed test), duration of (t(8) =-0.0983, p = 0.9241, two-tailed test) fLOM and total time spent in LOM (t(8) =- 0.6317, p = 0.5452, two-tailed test) for the same 3.0g/Kg I.P. Injection of ethanol (Fig. S3). Representative traces of mice activity showed that mice with Ca^v^3.1 KD in MD (Fig. S4- A) had a more delayed and fragmented early period of LOM compared to MD LV- shControl and VB injected mice, as in mutant mice.

**Fig. 2:**
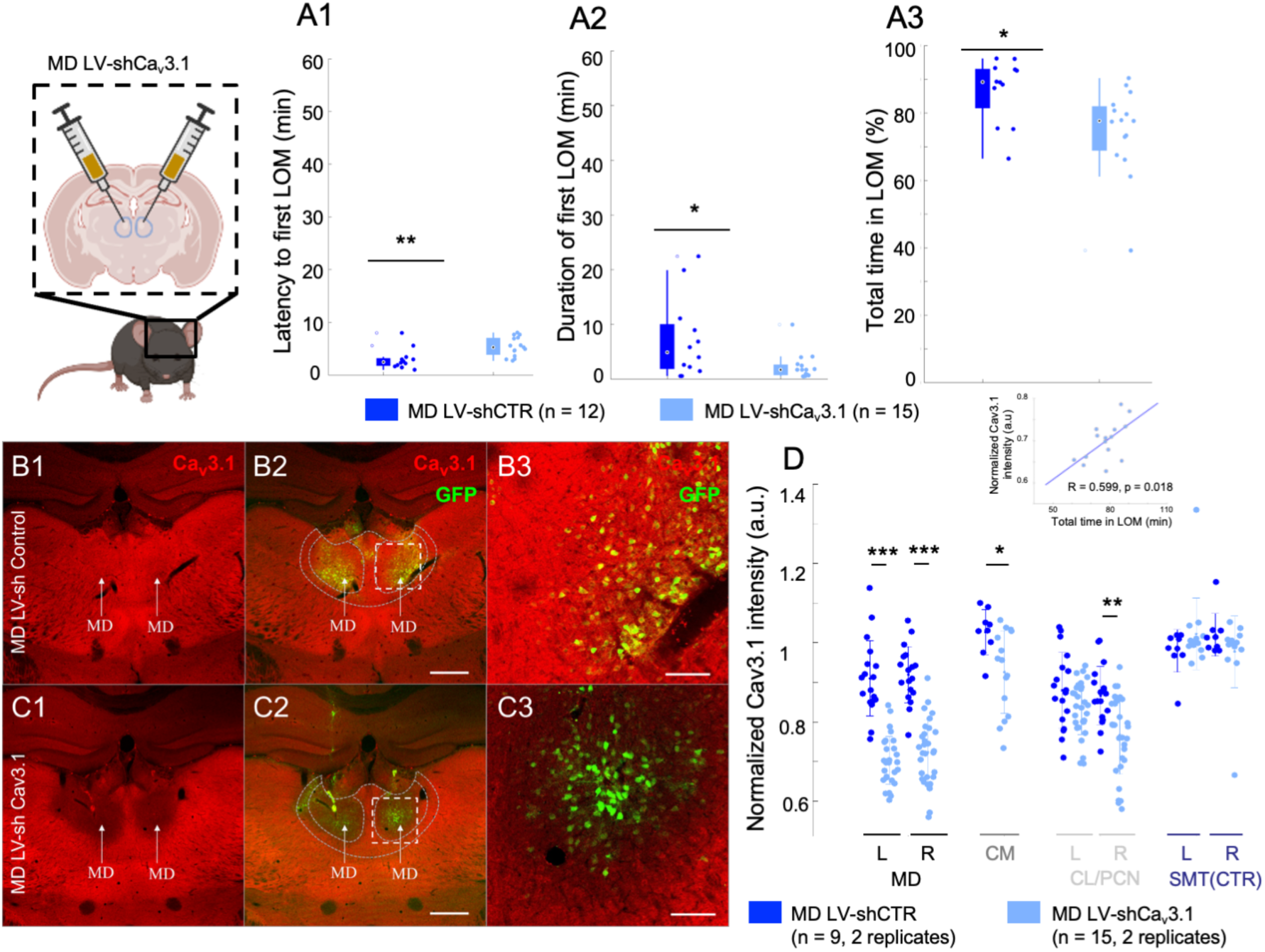
**Cav3.1 knock-down in MD increased ethanol resistance in mice.** (A) Quantification for the Latency to first LOM (fLOM; A1), the duration of the first LOM (A2) and the total time spent in LOM state (A3) over a recording duration of 60 min post I.P. injection of ethanol (3.0g/Kg) in lentivirus-shControl and shCa^v^3.1 knock-down mice for MD ; data is represented as a boxplot with individual mice shown as a scatter plot. * and ** indicates p<0.05 and p<0.01, respectively. (B) Representative brain coronal section stained using Ca^v^3.1 antibody (B1) showing the endogenous thalamic expression of Ca^v^3.1 in the MD of lentivirus(LV)-shControl injected mice; Ca^v^3.1 and GFP merging (B2) and higher magnification of the white dashed square in B2 (B3). Scale bars in (B2) and (B3) indicate 500 μm and 100 μm, respectively. (C) Representative brain coronal section showing the reduced Cav3.1 expression in the MD of LV-shCa^v^3.1 injected mice (C1); Ca^v^3.1 and GFP merging (C2) and higher magnification of the white dashed square in C2 (C3). Scale bars in (C2) and (C3) indicate 500 μm and 100 μm, respectively. (D) Normalized Cav3.1 intensity estimated for the nuclei MD, CM (centromedial), CL/PCN (Centrolateral/paracentral) and SMT (submedial thalamic nucleus). The quantification was performed as intensity per area for 2 replicates per side per mouse. *, ** and *** indicates p<0.05, p<0.01 and p<0.001, respectively (2 sample t-test). The data is shown as a scatter plot for all values and superposed with the mean and standard deviation error bars. Inset: We noted a positive correlation between the total LOM duration and the Cav3.1 intensity in MD (R = 0.599, p = 0.018).

We then characterized the change in Cav3.1 expression following the shControl and shCav3.1 knock-down injections in three test regions MD (left and right), CM (centromedial nucleus) and CL (centrolateral nuclei, left and right side) and a negative control region SMT (submedial thalamic nuclei, left and right side). The average intensity was obtained from two coronal brain slices for each mice used in the experiment (see Methods sections, Cav3.1 Intensity quantification). Our results show that the targeting of the knock-down was very specific to the bilateral MD (p<0.001; Fig. 2D). We noted that the CM (p<0.05) and a marginal unilateral knock-down of the CL were also observed (p<0.01). Notably, we tested the correlation between the level of knock-down in MD and the total time in LOM and observed a significant association (Fig. 2D inset; R = 0.599, p = 0.018). This result highlights that the Cav3.1 knock-down was specific to MD and with an intensity associated with ethanol-induced loss of motion.

During the open field test, Ca^v^3.1 null mutant mice showed significantly increased locomotor activity compared to WT mice as shown by total distance moved (Fig. S4-C; analysis of variance: GROUP F(3) = 8.45, p = 0.0004; Ca^v^3.1+/+ vs Ca^v^3.1-/-: p = 0.0001). The mice with MD-specific Ca^v^3.1 KD, however, did not show any significant difference in total distance moved compared to shControl-injected control mice (MD LV-shControl vs MD LV-shCa^v^3.1: p = 0.868; Holm-Sidak correction), indicating that the ethanol resistance in MD Ca^v^3.1 KD mice was not attributed by hyperlocomotion observed in Ca^v^3.1 KO mice.

### Lack of Ca3.1 in MD neurons removes thalamic burst in NREM sleep

Thalamic neurons are known to follow a state-dependent activity ^38^ ; however, the nature of this state-dependent activity has not been studied for the MD. In order to understand the relationship between MD neuron firing and level of consciousness, we investigated the association between the neural activity in MD and brain states at different levels of consciousness (Fig. 3 and S5).

**Fig. 3:**
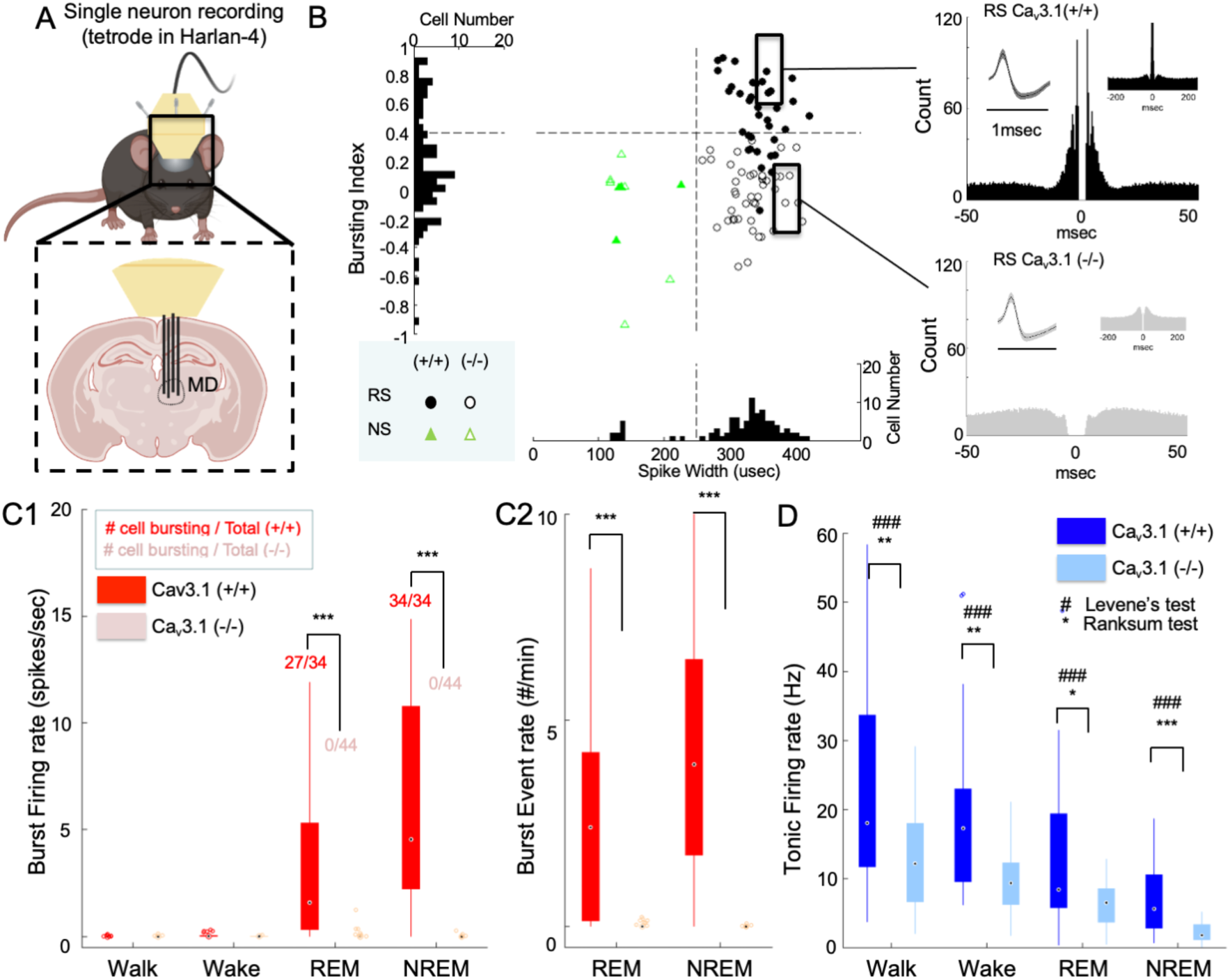
**Lack of Cav3.1 removed burst firing, and reduces neural activity and its variability in MD across natural conscious and unconscious states.** (A) Mice were implanted unilaterally with 4 tetrode wires to record single unit activity in the MD while in home cage (wake, sleep: NREM, REM) and forced walking task under ethanol (walk). (B - Left panel) Scatter distribution of spike width vs bursting index of MD regular spiking (RS, round shape) and narrow spiking (NS, triangle shape) neurons. Ca^v^3.1 (+/+) and Ca^v^3.1 (-/-) neurons are marked as filled and empty shapes, respectively . The histogram of the pooled Ca^v^3.1 (+/+) and Ca^v^3.1 (-/-) distribution is projected on each axis. (B - Right panel) Representative auto-cross correlograms of a RS neuron showing the presence and absence of fast spiking interval (burst firing) in Ca^v^3.1 wild-type and mutant mice, respectively. (C) Boxplots of Ca^v^3.1 wild and mutant burst firing rate (spikes/sec; burst spikes-only averaged over a state duration) (C1) and burst event rate (#/min; number of burst event averaged over a state duration; see burst definition in methods section) (C2) in RS neurons of the MD during NREM sleep, a stage known for the presence of bursting firing mode in thalamic neurons, for Ca^v^3.1 (+/+) and Ca^v^3.1 (-/-) mice; The inset numbers in C1 indicate the number of neurons showing burst firing (more than 1 event in 10 min) over the total number of single neurons identified. (D) Boxplots of Ca^v^3.1 wild and mutant tonic firing rate (spikes/sec) in RS neurons of the MD during walking (FWT), wake (home cage), REM and NREM sleep (home cage) for Ca^v^3.1 (+/+) and Ca^v^3.1 (-/-) mice. Group and brain state effect and interaction were assessed using a two-way repeated-measure ANOVA. For post-hoc, two samples Ranksum test comparison *, ** and *** indicate p-value lower than 0.05, 0.01 and 0.001, respectively. For two sample Levene’s test for homoscedasticity #, ## and ### indicate p-value lower than 0.05, 0.01 and 0.001, respectively. Pearson Ranksum correlations between brain state and total firing for Ca^v^3.1(+/+) and Ca^v^3.1(-/-) is indicated above boxplots.

We observed two major populations of neuronal spike waveform present in MD single unit recordings of Ca^v^3.1 (+/+) mice, also described in previous works ^39,40^ : 1) a majority of regular spiking (RS) cells characterized by wide spike waveform (36/39 neurons, 92.3%), i.e. spike-to-valley width >250 μsec (Fig. 3B) and high bursting propensity; 2) a minority of narrow spiking (NS; also known as fast spiking) cells showing short spike-to- valley width, i.e. <250 μsec, lower bursting characteristics (Fig. 3 and S6-A3 and -A4) and fast-paced tonic firing (10-50Hz; data not shown). The RS and NS neurons were found in MD of WT and mutant mice; however, MD RS mutant neurons showed an absence of short inter-spike intervals (ISI, i.e. indicative of the absence of burst) in auto cross- correlogram (Fig. S6) and a clear reduction in total bursting represented as bursting index (Fig. 3-B; ratio of spikes count <10 ms and >50 ms based on auto-cross-correlogram). Since RS cells have the profile of the major population of MD, i.e. excitatory neurons, we focused on the analysis of RS neurons mainly in the remainder of this study.

During the deep sleep state non-rapid eye movement (NREM), thalamic neural firing is known to switch from tonic to burst firing ^41^. We found that a lack of Ca^v^3.1 T-type calcium channels resulted in a near absence of burst (see Methods for definition) in mutant mice (Fig. 3-C1 and C2; 4/44 bursting neurons; Z(77) = 7.20, p < 0.0001, Ranksum test) compared to WT mice (34/34 bursting neurons; 5.76 ± 5.51 burst events/min).

### Lack of Ca3.1 reduces neuronal activity across all brain states in MD

In addition, we observed that the mutant mice showed a significant lower total firing rate (main factor group: F(1,186) = 16.5, p = 0.0001; interaction group x brain state: F(3,186) = 4.72, p = 0.0034) and a reduced variability (p <0.0001 for all brain states; Levene’s test) in most brain states compared to the wild type mice, indicating that the lack of Ca^v^3.1 t- type channels results in an overall reduction in neural activity in RS neurons.

RS neurons of MD in Ca^v^3.1 wild type and mutant mice showed a significant change in overall firing across walking, waking (home cage), NREM and REM sleep states as shown by a repeated measure ANOVA (Fig. 3-D; main factor brain state: F(3,186) = 104.96, p <0.0001). In addition, Ca^v^3.1 wild type (R = -0.534, p = 1.6e-10, Spearman ranked correlation) and mutant (R = -0.689, p = 6.8e-20, Spearman ranked correlation) showed a significant negative correlation between neural firing and brain state. Assuming an ordering from higher to lower state of consciousness, these results indicate that MD firing is associated with level of consciousness independently from the Ca^v^3.1 T-type channels in wild type and mutant mice. Importantly, this result indicates that Ca^v^3.1 t-type calcium channels are critical excitatory ion channels that control the overall neural activity along with the brain state. In other words, mutant mice exhibit a less clear distinction in the neural activities associated with wakeful and unconscious states.

### Under ethanol, MD neurons lacking Ca^v^3.1 show no burst and a wake state-like neural activity

In order to identify the mechanism linked to Ca^v^3.1 mutant mice ethanol-resistant phenotype, we recorded neural firing of neurons during the FWT and following a hypnotic dose of ethanol (3.0 g/kg, I.P. injection). We focused on the first loss of motion (fLOM) as it is most analogous to the classical LORR and showed the most consistency between animals ^36^. fLOM also illustrates best the acute effect induced by ethanol before secondary metabolization enters in play.

Under ethanol, we observed that in WT mice a majority of neurons showed burst firing mode (Fig. 4-A1; 20/33 bursting neurons). We found a significant higher burst event rate (Fig. 4-B1; p <0.0001, Ranksum test with Holm-Bonferroni correction) and in the ratio of burst-to-total spike (Fig. 4-B2; p <0.0001, Ranksum test with Holm-Bonferroni correction) comparing walk (awake active) to fLOM (unconscious, unresponsive). Mutant neurons, consistently with NREM data, did not show burst firing during fLOM (0/36 bursting neurons).

**Fig. 4:**
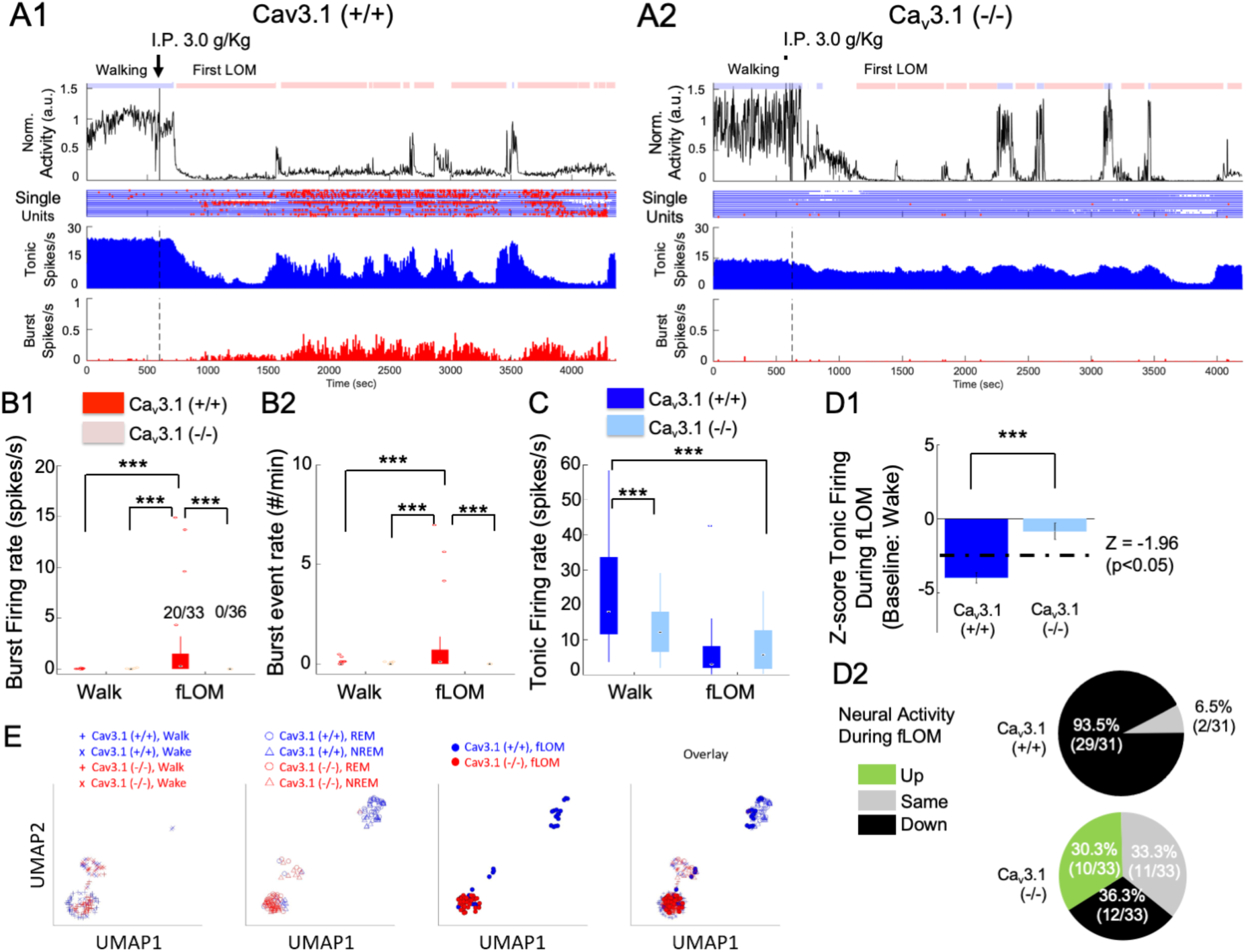
**Resistance to the loss of consciousness in Ca^v^3.1 mutant is associated with maintenance of neural activity and absence of burst.** (A) Representative time plot for, from top to bottom, the normalized activity, single unit raster plot (blue and red dots for tonic and burst firing, respectively), population mean firing (spikes/sec) and burst-to-total spike ratio (%) for Ca^v^3.1 (+/+) (A1) and Ca^v^3.1 (-/-) mice (A2). (B) Boxplots of Cav3.1 wild and mutant burst firing rate (spikes/sec)(B1) and burst event rate (#/min)(B2) in RS neurons of the MD during FWT walk (pre I.P. injection) and during FWT first loss of motion (fLOM, post I.P. injection) for Ca^v^3.1 (+/+) and Ca^v^3.1 (-/-) mice; The inset numbers in B1 indicate the ratio of the number of neurons showing burst firing over total neurons. Multiple comparisons were performed using two samples Ranksum test or paired sign rank test with Holm-Bonferroni correction. *** indicates a p-value < 0.001. (C) Boxplots tonic firing rate (spikes/sec) in RS neurons of the MD during FWT walk (pre I.P. injection) and during FWT first loss of motion (fLOM, post I.P. injection) for Cav3.1 (+/+) and Ca^v^3.1 (-/-) mice. Group and brain state effect and interaction were assessed using a two-way repeated-measure ANOVA. Post-hoc multiple comparison performed using two samples rank sum test or paired sign rank test with Holm-Bonferroni correction.*** indicates a p-value < 0.001. (D) Normalized Z-score firing during fLOM with respect to wake state (home cage) firing (D1). WT and mutant distribution and cell count based on fLOM Z-score showing increase (>1.96), no change (<1.96 and >-1.96) or decrease (<-1.96) in firing in Ca^v^3.1 (+/+) and Ca^v^3.1 (-/-) mice (D2). UMAP (uniform manifold approximation and projection) 2-dimensional representation of wakeful states (walk: + symbol; wake: x symbol; left panel), sleep states (REM: empty triangle symbol; NREM: empty round symbol; middle panel) and fLOM state (filled round symbol; right panel) of Ca^v^3.1 (+/+) (blue symbols) and Ca^v^3.1 (-/-) (red symbols) mice. The all-state overlay is depicted on the far right panel.

Notably, in WT, we observed that ethanol induced a significant decrease in total firing from walking to fLOM states (Fig.4-A1 and -C; p <0.0001, Rank Sum test with Holm-Bonferroni correction) and well below wakefulness level (Home cage awake state). As in sleep, we found that a majority of RS neurons showed decreased tonic firing (total number of spikes) together with an increase in burst firing, indicating a switch in firing mode under ethanol sleep. Interestingly, the mutant mice did not show a significant decrease in total firing (Fig. 4-C; p = 0.130, Ranksum test with Holm-Bonferroni correction) and showed no burst as in sleep.

We quantified the change in activity of individual neurons using Z-score normalized to the home cage wakeful state. Here, we also observed that WT RS neurons showed a significantly reduced Z-score under ethanol fLOM whereas in mutant mice cells did not (Fig. 4-D1; normalized from home cage wake state; t(62) = -5.1400 , p < 0.0001, student t-test). Remarkably, we found that a majority of WT MD neurons (29/31) showed individual significantly decreased Z-scores (Fig. 4-D2; z-threshold defined from a p-value of 0.05). During fLOM, mutant RS neurons subdivided into three populations (Fig. 4-D2) with decreasing (12/33, 36.4%), maintaining (11/33, 30%) and increasing (10/33, 30.3%) activity as measured by the Z-score with respect to wakefulness (i.e. home cage wake state). These results were consistent in individual mice (Fig. S7A) and the distribution of neural population spiking (Fig. S7B), validating that significant drop in neural activity is associated with loss of movement.

Finally, we asked whether the firing modes and properties (tonic firing rate, burst firing rate; see supplementary methods) of single MD neurons would form distinct qualitative representation of “brain stages” using a lowered dimensional UMAP representation (Uniform Manifold Approximation and Projection^42^ ). We observed that for awake and active (i.e. walk), the brain state representation formed two adjacent clusters that confounded both wild and mutant neurons (Fig. 4E, left panel). The REM and NREM states, the wild type neurons formed 2 additional interconnected clusters, whereas the mutant neurons tend to overlap with the clusters attributed to the “awake” brain state (Fig. 4E, second to left panel). Ethanol induced fLOM, similarly to REM and NREM clusters, was distinct from awake clusters in wild type mice and overlapped with the NREM clusters (Fig. 4E, third to left panel). Here also, mutant MD neurons showed overlap with the awake clusters rather than the “low consciousness” brain states. These results indicate that the firing mode and properties could define a brain state representation that shows distinctions in levels of consciousness. Moreover, the mutant showed a representation of “low consciousness” states overlapping with wild type “awake” states consistent with the hypothesis of resistance to loss of consciousness.

Altogether, these results indicate that, in WT, ethanol induced a strong reduction in neural activity and a switch to bursting firing mode correlated with loss of consciousness. However, under ethanol, MD mutant neurons maintained their activity to a level within home cage wake state without switching to bursting. This indicates that the drop in neural activity under ethanol is modulated by Ca^v^3.1 t-type calcium channels. In its absence, MD mutant neurons display an overall reduced activity in all brain states, however, under ethanol, remain within state wakeful levels.

### Under ethanol, 20 Hz neurostimulation of MD induces mutant-like resistance to loss of consciousness in WT mice

We observed that the maintenance of neural activity in MD excitatory neurons might be at the origin of the ethanol resistance in mutant mice. We hypothesized that artificially maintaining MD neural activity within the wakeful level would sustain consciousness under ethanol. In addition, we hypothesized that the triggering of burst firing under ethanol would potentiate loss of consciousness under ethanol. To test this possibility, we used electric and optogenetic stimulations during the FWT in WT mice under a hypnotic dose of ethanol.

MD neurons in WT mice showed a spike firing range 0-50 Hz with an average neural firing around ∼20 Hz during home cage wakefulness (Fig. S7-B). Using the 20 Hz stimulation (Fig. 5-A, inter-pulse interval = 50 msec, pulse width = 6.25 msec) in MD neurons transduced with excitatory channelrhodopsin ^43,44^ (aav-syn-ChR2-sfGFP; Fig. 5-B and Fig. S9; see Methods), we observed an increase in ethanol resistance, which was demonstrated by a significant increase in latency to fLOM (Fig. 5-D1; Z(13) = -2.372, p = 0.013; Ranksum test). The duration of fLOM (Fig. 5-D2; Z(13) = 2.256, p = 0.020; Ranksum test) and total time spent in LOM (Fig. 5-D3; Z(13) = 2.488, p = 0.009; Ranksum test) were also significantly reduced. We verified that optogenetic stimulation of MD neurons at 20Hz (Fig. S8) induced action potentials at the same frequency with a latency response of about ∼5 ms (Fig. S8-B2; pulse width = 6.25 msec). We also observed that, although marginally higher, optogenetic stimulation did not induce any significant increase in locomotor activity (Fig. S8-C; F(1,12) = 3.6232, p = 0.0812) in the control and stimulated group which indicates that the stimulation-induced increase in ethanol resistance was not due to an increase in locomotor activity.

**Fig. 5:**
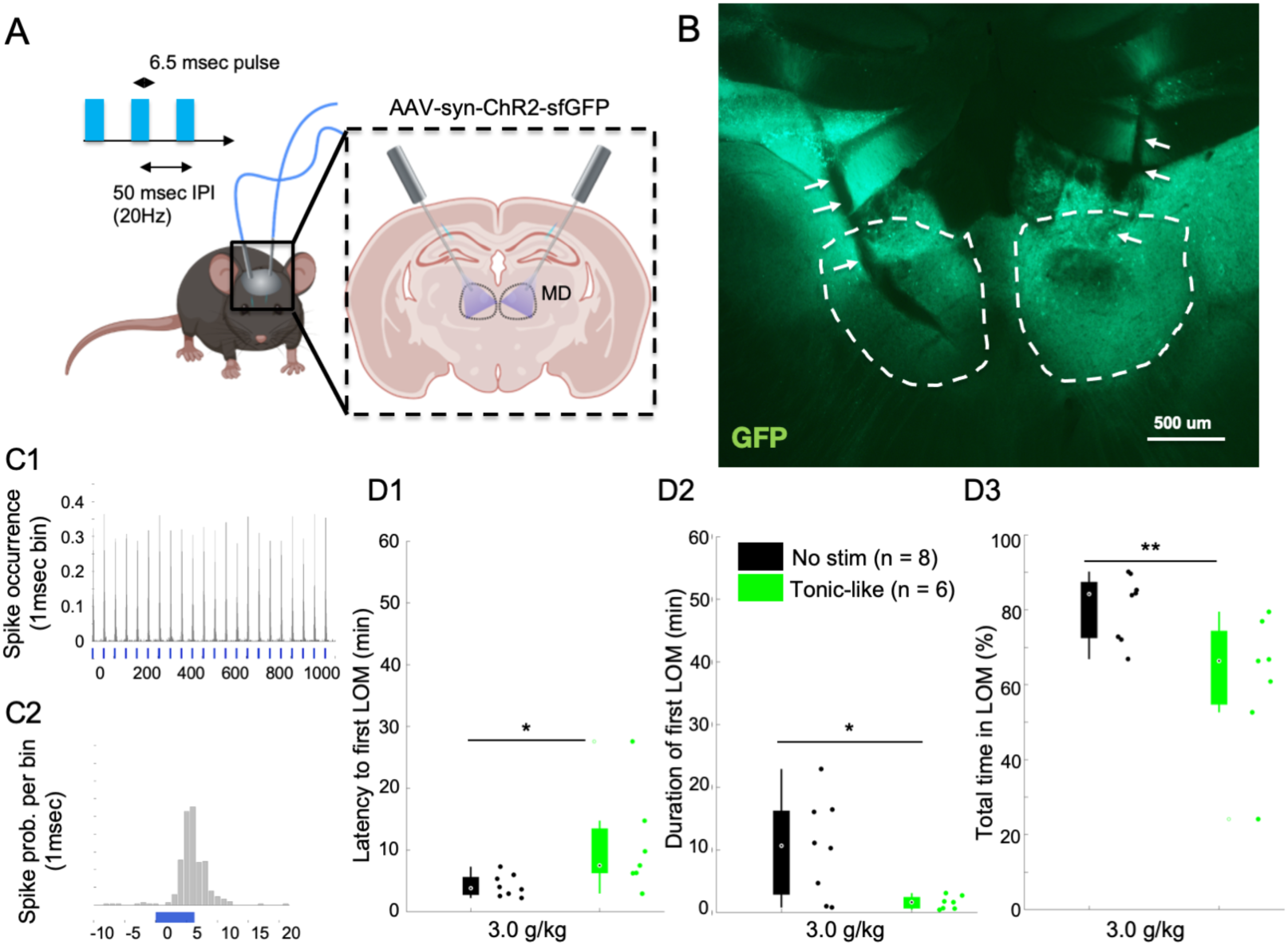
**Optogenetic 20 Hz stimulation of MD in WT mice mimics ethanol resistance.** (A) Mice were transduced bilaterally in the MD with an AAV-syn-CHR2-sfGFP and implanted with bilateral optic fibers targeting MD with an entry angle of 30 deg relative to the sagittal plane. We used a stimulation protocol of tonic-like pulses at 20 Hz with 50 ms inter-pulse-interval (IPI) and 6.25 ms pulse duration. (B) Representative expression of ChR2-sfGFP in the MD (dashed white lines) with fiber optic ending (white arrows). (C) In vivo response of a MD RS neurons to the 20Hz tonic stimulation protocol using 6.25 ms pulse at 20Hz and laser power of 6.0 mW (C1); magnification of the peristimulus response of the neuron around the laser pulse (1 ms bin; C2). Latency to first LOM (D1), duration of the first LOM (D2) and total time spent in LOM state (D3) over a recording duration of 1 hour post I.P. injection of 3.0g/Kg of ethanol are shown for the control group (No Stim) and stimulated group (Tonic-like). * is for p<0.05, ** is for p<0.01, *** is for p<0.001.

In order to validate this observation, we then bilaterally implanted mice with twisted wires for bipolar, local electric stimulation of MD (Fig. S11). As in the optogenetics experiment, we used a continuous pulse train of 20Hz electric stimulation (Fig. S10-A; upper panel; inter-pulse interval = 50 msec, pulse width = 1 msec) in addition to a burst-like stimulation (Fig. S10-A; lower pane; 4 pulses at 4 ms interval and interburst interval of 1 sec) that showed, respectively, tonic-like and burst-like entrainment in thalamic neurons ^45^. We observed that our 20Hz tonic-like stimulation significantly increased the latency to fLOM (Fig. S10-B1; tonic-like vs no stim.: p = 0.008; tonic-like vs burst-like: p = 0.007; Ranksum test with Holm-Bonferroni correction) and significantly decreased the total time spent in LOM (Fig. 5-B3; tonic-like vs no stim.: p = 0.008; tonic-like vs burst-like: p = 0.007, Ranksum test with Holm-Bonferroni correction). No significant changes in the duration of the first LOM were observed (duration of fLOM (Fig. 5-B2; tonic-like vs no stim.: p = 0.917; tonic-like vs burst-like: p = 0.606; Ranksum test with Holm-Bonferroni correction).

Interestingly, we observed that burst-like electrical stimulation (Fig. S10-A; lower panel; inter-spike interval = 4 msec, inter-burst interval = 1 sec, pulse width = 1 msec) did not induce any significant change in ethanol sensitivity compared to the no stimulation group (Latency to fLOM: p = 0.365; duration of fLOM: p = 0.835; total time spent in LOM: p = 1; Ranksum test with Holm-Bonferroni correction). This result suggests that burst firing alone might not have a role in ethanol resistance.

Altogether, these results suggest that the maintenance of MD firing at wakefulness level (20Hz) causally drives resistance to loss of consciousness after a hypnotic dose of ethanol. Burst-like stimulation alone did not promote or reduce loss of consciousness. This result supports the idea that neural activity maintenance in MD promotes the maintenance of consciousness even under heavy sedatives.

## DISCUSSION

In this work, we identified that the neural activity in MD plays a causal role in the maintenance of consciousness. Whole body Ca^v^3.1 KO and MD-specific Ca^v^3.1 KD mice showed resistance to loss of consciousness induced by hypnotic dose of ethanol. In WT mice, MD neurons demonstrated a reduced firing rate in natural (sleep) and ethanol-induced unconscious states compared to awake states. This neural activity reduction was impaired in KO mice. In particular, transition to an unconscious state was accompanied with a switch of firing mode from tonic firing to burst firing in WT mice whereas this mode-shift disappeared in KO mice. Finally, optogenetic or electric stimulations of the MD after ethanol injection were sufficient to induce a resistance to loss of motion, supporting that the level of neural firing in the MD is critical to maintain conscious state and delay unconscious state. We showed that the expression of Ca^v^3.1 t-type calcium channels in MD is a cellular modulator associated with this effect.

### MD is a modulator of consciousness

The role of mediodorsal (MD) thalamic nucleus in perception, attention ^46^ and emotional control ^45,47^ has been the dominant focus thus far. The recent investigations on thalamic control of consciousness revealed that nuclei within dMT holds an important modulatory function in the interaction of attention and arousal ^9,30^. Particularly, centromedian thalamic nucleus, and not VB, showed rapid shifts in LFP preceding brain state transitions such as NREM and propofol-induced anesthesia ^8^. The centrolateral thalamic nucleus was implicated in the modulation of arousal and improvement of consciousness during seizure^32^ and paraventricular thalamic nucleus showed critical involvement in wake/sleep cycle regulation ^48^ . The mediodorsal thalamic nucleus, however, has rarely been included as a possible pathway in the direct modulation of consciousness ^33^. The MD receives projections from various parts of the basal forebrain ^9^ and brainstem nuclei such as the pedunculopontine nucleus that control the ascending pathway of arousal and attention ^49^. The MD is known to innervate limbic region, basal ganglia and medial prefrontal cortex ^50^ and increased activity in MD might modulate the stability of cortical UP states (e.g. awaken, aroused and attentive states) and synchronization ^9,26^. Thus, MD might be a major hub involved in cortical state control and brain state stabilization.

Supporting the brain state stabilization theory and the ethanol resistance of Cav3.1 mutants, Choi et al.^34^ demonstrated that the loss of Cav3.1 T-type calcium channel reduced the bilateral coherence between PFC and MD under ketamine anesthesia and ethanol hypnosis, especially in the delta frequency bands. More importantly, under propofol anesthesia, Bastos et al.^35^ showed that intralaminar nucleus and MD stimulation lead to increased wake-up subscore and arousal, together with an increased in cortico-cortico and thalamo-cortical slow (delta) frequency power.

In the present study, we observed that MD KD (Fig. 2A), but not VB KD (Fig. S3) of Ca^v^3.1 increased and is associated (Fig. 2D) with ethanol resistance in mice. We found that MD neurons in Ca^v^3.1 mutant mice exhibited tonic firing within range of wakefulness (Fig. 3 and 4), indicative of resistance to ethanol and wake-like brain state. In addition, we found a strong association between the normalized tonic firing in MD and the arousal through brain states (i.e. walk to wake to sleep states), supporting that MD tonic firing could be interpreted both as a thalamic readout and a modulator of the brain state ^11^ (Fig. 3). Finally, direct optogenetic and electric MD stimulation increased resistance to loss of consciousness in WT mice (Fig.5 and Fig. S10). To our knowledge, this is the first report demonstrating the causal involvement of mediodorsal thalamic nucleus in the modulation of wakefulness and the resistance to ethanol-induced loss of consciousness in mice.

### Ca^v^3.1 T-type calcium channels drive thalamic firing mode and activity

The decrease of absolute firing rate observed in thalamic neurons of Ca^v^3.1 mutant mice supports the polyvalent role of Ca^v^3.1 in controlling both burst and tonic firing in the thalamus. Ca^v^3.1 channels are major contributors of excitability, and in their absence or blockade, leads to a reduced neural excitability and stability and lower tonic relay of thalamocortical cells under wake-like state ^51,52^. The burst and tonic firing-mediated response of thalamic neurons under sensory stimulation and under the control of thalamocortical layer 6 projecting neurons was found to recruit Ca^v^3.1 t-type calcium channels to differentiate salient novel stimuli versus complex coded information ^53^. Therefore, the nonlinear amplification and regularization of excitatory postsynaptic potentials (EPSPs) by Ca^v^3.1 T-type calcium channels through complexes such as with metabotropic-glutamatergic receptor 1 (mGluR1) ^54^ or the role of a “T window” ^55^ would explain how the lack of T-current in mutant mice could result in an overall reduced excitation of thalamic neurons. Ca^v^3.1 T-type is therefore a major excitatory ion channel of the central thalamic neurons.

### The lower variability in MD Firing reflects Ethanol Resistance in Ca^v^3.1 mutant mice

Under acute hypnotic dose of ethanol, two mechanisms might favor the reduction in firing in MD: 1) an increase in synaptic and extra synaptic GABAergic inhibition ^17^ and/or 2) reduced NMDA synaptic transmission ^7^. The presence of burst firing during fLOM, and during LOM in general, supports that MD neurons might have been subject to GABA receptor-mediated hyperpolarization, a necessary condition for the de-inactivation of Ca^v^3.1 T-type burst. However, considering the dramatic difference in tonic firing observed during the FWT following I.P. injection of ethanol, the change in tonic firing in MD was the focus of our analysis.

We observed a reduction in neural firing under ethanol sleep conditions in WT mice (Fig. 4-C, -D1 and -D2) suggesting that low firing level should be associated with a state of low consciousness as observed during NREM sleep. Mutant RS neurons in MD showed an overall lower excitability and variability of firing in various natural conscious and unconscious states compared to wild type mice. Remarkably, Ca^v^3.1 mutant mice exhibited a clear increased locomotor activity and an increased resistance to ethanol. The general lower firing rate and the high “arousal” observed in mutant mice suggests that the relative change from state to state in tonic firing in MD, and not the absolute value of firing, might be a better correlate of change in brain state in the mice. Our optogenetic and electrical stimulation showed that a sustained tonic-like stimulation in the MD at 20Hz (Fig. 5-A), a physiological relevant firing rate in wake state (Fig. 4), could increase ethanol resistance in WT mice. Reducing MD firing using phasic inhibition under ethanol, potentially leading to inhibition and rebound burst ^56^, could also increase the duration of the fLOM in WT mice injected with a lower dose of ethanol (Figure S12; 2.0 g/kg). We propose that the relative change in firing rate in MD RS neurons might be an important driver and indicator of the change of transition in and out of consciousness as demonstrated for other nuclei of the dMT ^8,9,32,48^. Therefore, the low variability in firing of MD in Ca^v^3.1 mutant mice might be the driving force for the higher resistance to loss of motion under ethanol. In mutants, brain states might be less distinguishable leading to frequent sleep stages switch^23^ or resistance to unconsciousness^34^.

### Ca^v^3.1 T-type Calcium and Burst during Low Conscious State

Burst, as a result of Ca^v^3.1 T-type calcium channel de-inactivation/activation, is thought to control the gating of sensory-motor stimuli ^57,58^ and modulate attention towards novel stimuli rather than the transmission of details ^53,57,59^. Previous reports highlighted the importance of burst in the stabilization of low level of consciousness ^13,22,23^ suggesting a direct role for burst, while no mention of the importance of tonic firing was made. We found that the propensity for burst during ethanol-induced LOM (Fig. 4-A1, -B1 and -B2; fLOM: 20/33 bursting neurons; 0.79±1.63 Burst event/min) was lower than in NREM (NREM: 34/34 bursting neurons; 5.76 ± 5.51 burst events/min) and higher than during wakefulness (Wake: 5/36 bursting neurons; 0.16±0.31 Burst event/min). In addition, burst-like electrical stimulation of MD did not significantly affect ethanol resistance (Fig. S10). Although burst-like stimulations are highly artificial and do not recruit T-current and associated mechanisms following low-threshold burst, they allow for the reproduction of the influence of TC burst firing on target centers^45^, including thalamo-cortical and thalamo-thalamic efferents.

Interestingly, under lower doses of ethanol (I.P. injection of 2.0g/kg of ethanol) mutant and WT alike showed similar levels of resistance to ethanol. We observed a phasic inhibition of MD neurons in WT, capable of inducing partial silencing ^60^ and rebound bursts ^61^ (Fig. S12; 1sec ON-OFF using archaerhodopsin-mediated inhibition), did increase fLOM duration mostly (Z(13) = -2.214, p = 0.022, Ranksum test). This result supports that in the context of hypnotic dose of ethanol, the apparition of burst might correlate with unconscious state stability rather than induction. Burst stimulation without inhibition did not have this effect (Fig. S10 and S11). Currently, our data does not allow us to formulate any clear conclusion on the direct role of burst events during fLOM. We propose that the absence of burst and an accompanying effect of maintenance of tonic firing under ethanol in MD was responsible for the observed increase in resistance and maintenance of activity in Ca^v^3.1 mutant mice.

### A bidirectional modulation of Ca^v^3.1 expression and alcoholism

In human mutation of Ca^v^3.1 T-type channels exhibit mental disorders including cerebellar ataxia, absence seizure, schizophrenia and autism^62^. Remarkably, mutation in voltage-gated calcium channels, including Ca^v^3.1 leads to ethanol resistance and alcohol-seeking behavior ^21^. Reversibly, chronic exposure to ethanol intake is known to impair sleep ^63^ and increase ethanol resistance. Previous studies have found an alteration in T-type calcium channel expression following chronic exposure to ethanol in non-human primates ^64^ suggesting that a reduced T-current and the resulting sustained thalamic tonic firing could be a possible mechanism for early stage ethanol resistance in alcoholic subjects, which increases the conversion probability from casual to compulsive consumption of ethanol. The lack of burst and sustained tonic firing might impair the stabilization of sleep, and in turn chronic sleep impairments might engage addiction related networks. Mechanisms such as adenosine receptor depreciation ^65,66^ or GABA-receptor potentiation ^18,67,68^ would enhance ethanol resistance and addiction ^69,70^ spiraling into further sleep fragmentation, memory consolidation deficit, impulsivity and other impairments associated with alcoholism.

## ACKNOWLEDGMENTS

This work was supported by the grant IBS-R001-D1 and IBS-R001-D2 from the Institute for Basic Science, Korea. We would like to thank Dr. Gireesh Gangadharan for his precious help in the editing of this manuscript.

## Author Contributions

C.-F. V. L., S.K. and J.-H. L. performed the experiments.

C.-F. V. L. and J.-H. L. performed the data analysis.

J.-H. L. and S. K. performed the histological work.

1. J. K. provided the aav9-syn-ChR2-sfGFP virus for the optogenetic study.

C.-F. V. L, J.-H. L., S.K. and H.S. designed the study and wrote the manuscript.

## Data Sharing

Part of the analyzed data and codes are available on the open access platform, mendeley:

Latchoumane, Charles-francois (2024), “Mediodorsal thalamic nucleus mediates resistance to ethanol through Cav3.1 T-type Ca2+ regulation of neural activity”, Mendeley Data, V1, doi: 10.17632/7fr427426m.1

Additional data (heavy recording and images) can be provided upon reasonable requests.

## METHODS

### Animals

Ca^v^3.1 heterozygous mice (Ca^v^3.1^+/−^) were maintained in two genetic backgrounds, 129/svjae and C57BL/6J. All experiments used Ca^v^3.1^-/−^ mice and their WT littermates in the F1 hybrid generated by mating CaV3.1+/− mice from these two genetic backgrounds. Mice were maintained with free access to food and water under a 12-h light/12-h dark cycle, with the light cycle beginning at 8:00 AM. Animal care was provided and all experiments were conducted in accordance with the ethical guidelines of the Institutional Animal Care and Use Committee of the Institute of Basic Science and the Korean Institute of Science and Technology. All experiments were conducted using 12- to 16-wk-old male mice.

### Surgery for electrophysiological recordings and neurostimulation

The surgical implantation of electrodes (EEG, EMG and/or tetrode Microdrive) and virus injection procedures were performed under 0.2% tribromoethanol (Avertin) anesthesia (20 mL/kg i.p.). Following anesthetic administration, mice (11-week-old for electrode implantation; 10 weeks for virus injection) were fixed in a stereotaxic device (David Kopf Instruments). For chronic recording of EEG and EMG, a stainless-steel screw electrode was fixed into the skull over the right parietal hemisphere and an uncoated stainless-steel wire was tied to the nuchal muscle, respectively. For in vivo freely moving single unit recording, we used a Harlan 4 Drive (Neuralynx inc.) mounted with 3 to 4 tetrode wires inserted to the caudal region of the right mediodorsal thalamic nucleus (anteroposterior, −1.4; lateral, +0.4; depth: 3.2 mm). Single tetrode wires were prepared from 4 twisted nichrome- formvar/PAC wires (Kanthal precision technology, OD 0.0127 mm) and gold plated to achieve an impedance range of 150-400kΩ (1kHz, in saline solution). A period of 7 days was given to allow a complete recovery from the surgical procedure. For all chronic implantation of electrodes, an additional screw was positioned over the occipital region and used as a reference.

### Optogenetic neurostimulation

For optogenetic experiment, 16 mice were bilaterally injected with aav9-syn-ChR2-sfGFP virus in the mediodorsal thalamus and implanted with optic fiber guides (125 μm core diameter, Doric Lenses inc.) positioned at 30 degree angle from the transverse plane. The mice were given 2∼3 weeks to recover and to allow for the viral expression. These mice were then randomly assigned to a no stimulation (n = 8) and a 20 Hz stimulation group (n = 6). The mice received the stimulation immediately after being placed in the treadmill, then received the i.p. injection of 3.0 g/kg of ethanol as in other experiments. We discarded 2 mice due to a low viral expression found after histological analysis.

In order to measure the neural response to optogenetic stimulation, we implanted one mouse unilaterally (right MD) with a Harlan 4 Drive (4 tetrodes) converging with a single optic fiber (right side, 30 deg inclination). This mouse received optogenetic stimulation in a home cage resting condition and at frequency 1-5-10-20-40 Hz with a fixed stimulation pulse of 6.5 ms (Fig. S8). Using these recordings, we verified the fidelity between the triggered laser stimulation and the single unit response in the vicinity of the laser illumination. The spike per stimulation trial, spike initiation success rate and the delayed triggered spiking (jittering) were estimated from these recordings. For all simulations, we used a high stability 473 nm (blue, MFB-III-473-AOM; Changchun New Industries Optoelectronics Technology Co., Ltd.) fiber coupled (FC) at an intensity of 6.0 mW. Laser triggering was performed using a Pulsepal pulse generator (gen1, open-source; https://open-ephys.org/pulsepal) or Master-8 (A.M.P. Instruments, Israel) pulse stimulator. **Electric neurostimulation.** For experiments using electrical stimulation, 18 mice were implanted with bilateral twisted dual stainless-steel wires (A-M systems, PFA coated, 50 um diameter) targeting MD (anteroposterior, −1.4; lateral, +/-0.4; depth: 3.2 mm; from bregma). The wires were minted on a custom made 4x1 pin header connector and cemented. As in the optogenetic experiment, the mice received the stimulation immediately after being placed in the treadmill, then received the i.p. injection of 3.0 g/kg of ethanol. The mice were randomly distributed into 3 groups, Sham no stimulation (n = 6), 20 Hz tonic stimulation (n = 5; 100 μsec pulse duration with inter pulse interval of 50 msec), and Burst stimulation (n = 7; 4x pulses of 100 μsec duration at 250 Hz; inter burst interval of 1 sec). All stimulation were performed in a bipolar configuration (twisted wire, bilateral implants) and biphasic pulse (100 μA, current stimulation) using a 2100 isolated pulse stimulator (A-M Systems, inc.).

### Virus Injection

WT mice (10-week-old) were placed in the stereotaxic device following 0.2% tribomoethanol anesthesia (20 mL/kg i.p.). Custom elongated (Sutter Instrument Co.) borosilicate pipette (ID: 0.05 mm, OD: 0.07mm, World Precision Instruments, inc.) was used to inject 0.2 to 0.5 μL of virus solution at a rate of 0.1μL/min (Hamilton syringe, pump) bilaterally in to the mediodorsal thalamic nuclei (anteroposterior, −1.4; lateral, +/- 0.4; depth: 3.2 mm). The injection pipette was then removed slowly after a diffusion period of 10min. A period of 2 to 3 weeks was given to allow viral infection to settle and a complete recovery from the surgical procedure.

### Ca^v^3.1 knock-down virus

For genetic knock-down of Ca^v^3.1 T-type calcium channels in the MD and VB in vivo, we used a lentivirus-mediated knockdown injection^43^. High-titer, concentrated lentiviral vectors (10^7^ TU/μl) expressing shCa^v^3.1(target sequence: 5’- CGGGAAGATCGTAGATAGCAAA-3’) or control shRNA (non-human or mouse shRNA : 5’-AATCGCATAGCGTATGCCGTT-3’) were prepared.

### Channelrhodopsin virus

Channelrhodopsin fused with superfolder GFP (ChR2-sfGFP) was designed and synthesized from published sequences using codon optimization for M. musculus (DNA 2.0). To express ChR2-sfGFP in the mouse brain, the AAV vector under a control of the human Synapsin promoter (aav-Syn) was generated using PCR-amplified human Synapsin promoter. Viruses were produced with Serotype 1 or DJ (Cell Biolabs, Inc.) and purified by CsCl gradients ^71^. The virus was injected at a volume of 0.5 μL in each side of the MD, followed by a bilateral implantation of optical fibers (100/125 μm, DP, Doric lens). The mice were given a period of 3 weeks to allow a strong expression of the channelrhodopsin channel following viral infection, as well as to recover from the surgical procedures.

### Cav3.1 Intensity quantification

For the quantification of Cav3.1 expression in the MD, we defined ROIs centered to the left and right of the MD (2x ROIs), CL/PCN (centrolateral, pericentral nucleus; 2x ROIs), and SMT (submedial thalamic nucleus; 1x ROIs; used as a control region of high Cav3.1 intensity, far from lentivirus injection). We added the CM (centromedial region; 1x ROI central only). All ROIs were predefined using a custom script in FIJI (ImageJ, doi:10.1038/nmeth.2019) and manually rectified to match anatomical position within the nuclei. We then run a custom MATLAB script to estimate the average intensity per area for all ROIs (11 ROIs defined in total per animal for each side, left and right). All intensities were then normalized to the average intensity of the SMT (highest expression region). We then compared the normalized Cav3.1 intensity for each animal for the factors side (left, right) and knock-down conditions (shRNA, shCAv3.1).

### Drugs

Tribromoethanol (Avertin) and Ethanol were purchased from Sigma-Aldrich. All drugs were administered by i.p. injection. The surgical implantation and virus injection procedures were performed under 0.2% tribromoethanol (Avertin) anesthesia (20 mL/kg i.p.). Ethanol injections were based from a prepared stock mixture of ethanol (26%) and saline and dosage were adjusted according to the experiments and the animal body weight (i.e. 2.0 g/Kg, 3.0 g/Kg and 4.0 g/Kg).

### Statistical Analysis

All statistical analyses were performed using Matlab and SPSS 17.0 (Statistical Package for the Social Sciences). Group differences were assessed using the Student t-test. In the case of low sample number (i.e. n<7) or distribution comparison of non-normal and/or non-equal variance number group difference were additionally confirmed using a nonparametric test (i.e. Wilcoxon Ranksum test/Signrank test). Multiple comparison p-value corrections were performed using a Holm-Bonferroni method. General longitudinal and group difference analysis were performed using repeated measures analysis of variance (ANOVA) and one/two-way ANOVA when advised. Linear correlation was performed using Pearson’s correlation coefficient.

### Mouse activity classification

Mouse activity was obtained using a video analysis and alternatively using an accelerometer placed on the head stage of the mouse when video wasn’t available. For video analysis, after histogram filtering of the mouse’s body color, the instantaneous activity was estimated as the frame-by-frame intensity difference followed by a 2D median filtering (3x3 pixel) and summed as the number of displaced pixels on camera. For the accelerometer, a zero-phase 5^th^ order Butterworth band-pass filter with cut off frequency of 0.5-20 Hz was used in order to remove the DC component; the instantaneous activity was derived as the root mean square (RMS) of x-, y- and z-axis filtered signals. The mean and standard deviation of the mean (STD) of the instantaneous activity was estimated in moving windows of 4sec duration (50% overlap).

The normalized activity index was obtained from the product of MEAN x STD (i.e. sustained activity and variability). Normalization was performed so that 1) complete cessation of activity approximate a value of 0 and 2) the 10 min walking baseline prior I.P. injection average a value of 1. A mouse was classified as not walking if its normalized activity was lower than the 95% confidence interval of baseline activity for a duration of at least 60 sec, and classified as walking otherwise. Non-walking states were reclassified as loss of motion (LOM) if the mouse’s normalized activity was maintained below 0.25 (lower quartile) for a duration of at least 30 sec. Adjustments were performed after manual video verification.

## REFERENCES

1. Alkire, M. T. & Miller, J. General anesthesia and the neural correlates of consciousness. In Progress in Brain Research (ed. Laureys, S.) vol. 150 229–597 (Elsevier, 2005).

2. Mashour, G. A. Top-down mechanisms of anesthetic-induced unconsciousness. Frontiers in Systems Neuroscience 8, (2014).

3. Morozova, T. V., Mackay, T. F. & Anholt, R. R. Genetics and genomics of alcohol sensitivity. Molecular Genetics and Genomics 289, 253–269 (2014).

4. Blomeley, C. P., Cains, S., Smith, R. & Bracci, E. Ethanol Affects Striatal Interneurons Directly and Projection Neurons Through a Reduction in Cholinergic Tone. Neuropsychopharmacology 36, 1033–1046 (2011).

5. Givens, B. S. & Breese, G. R. Electrophysiological Evidence that Ethanol Alters Function of Medial Septal Area Without Affecting Lateral Septal Function. J Pharmacol Exp Ther 253, 95–103 (1990).

6. Koob, Lewis, Meyer & Paul. Neuropharmacology of Ethanol: New Approaches. (Springer Science & Business Media, 2013).

7. White, G., Lovinger, D. M. & Weight, F. F. Ethanol inhibits NMDA-activated current but does not alter GABA-activated current in an isolated adult mammalian neuron. Brain Research 507, 332–336 (1990).

8. Baker, R. et al. Altered Activity in the Central Medial Thalamus Precedes Changes in the Neocortex during Transitions into Both Sleep and Propofol Anesthesia. J. Neurosci. 34, 13326–13335 (2014).

9. Schiff, N. D. Central Thalamic Contributions to Arousal Regulation and Neurological Disorders of Consciousness. Annals of the New York Academy of Sciences 1129, 105–118 (2008).

10. Llinás, R., Ribary, U., Contreras, D. & Pedroarena, C. The neuronal basis for consciousness. Philos Trans R Soc Lond B Biol Sci 353, 1841–1849 (1998).

11. Alkire, M. T., Hudetz, A. G. & Tononi, G. Consciousness and Anesthesia. Science 322, 876–880 (2008).

12. Steriade, M., McCormick, D. A. & Sejnowski, T. J. Thalamocortical oscillations in the sleeping and aroused brain. Science 262, 679–685 (1993).

13. Kim, D. et al. Lack of the Burst Firing of Thalamocortical Relay Neurons and Resistance to Absence Seizures in Mice Lacking α1G T-Type Ca2+ Channels. Neuron 31, 35–45 (2001).

14. Simms, B. A. & Zamponi, G. W. Neuronal Voltage-Gated Calcium Channels: Structure, Function, and Dysfunction. Neuron 82, 24–45 (2014).

15. Crunelli, V. et al. Dual function of thalamic low-vigilance state oscillations: rhythm-regulation and plasticity. Nat Rev Neurosci 19, 107–118 (2018).

16. Steriade, M., Nunez, A. & Amzica, F. Intracellular analysis of relations between the slow (< 1 Hz) neocortical oscillation and other sleep rhythms of the electroencephalogram. J. Neurosci. 13, 3266–3283 (1993).

17. Jia, F., Chandra, D., Homanics, G. E. & Harrison, N. L. Ethanol Modulates Synaptic and Extrasynaptic GABAA Receptors in the Thalamus. Journal of Pharmacology and Experimental Therapeutics 326, 475–482 (2008).

18. Jia, F., Pignataro, L. & Harrison, N. L. GABAA receptors in the thalamus: α4 subunit expression and alcohol sensitivity. Alcohol 41, 177–185 (2007).

19. Iyer, S. V. et al. α4-Containing GABAA Receptors are Required for Antagonism of Ethanol-Induced Motor Incoordination and Hypnosis by the Imidazobenzodiazepine Ro15- 4513. Front Pharmacol 2, (2011).

20. Newton, P. M. et al. A Blocker of N- and T-type Voltage-Gated Calcium Channels Attenuates Ethanol-Induced Intoxication, Place Preference, Self-Administration, and Reinstatement. J. Neurosci. 28, 11712–11719 (2008).

21. Shin, H.-S., et al. Mice lacking alpha 1G showing enhanced novelty-seeking and alcohol preference and therapeutic methods for mood disorders by modulating alpha 1G T-type calcium channels. (2005).

22. Anderson, M. P. et al. Thalamic Cav3. 1 T-type Ca2+ channel plays a crucial role in stabilizing sleep. Proceedings of the National Academy of Sciences of the United States of America 102, 1743–1748 (2005).

23. Lee, J., Kim, D. & Shin, H.-S. Lack of delta waves and sleep disturbances during non-rapid eye movement sleep in mice lacking α1G-subunit of T-type calcium channels. Proceedings of the National Academy of Sciences 101, 18195–18199 (2004).

24. Petrenko, A. B., Tsujita, M., Kohno, T., Sakimura, K. & Baba, H. Mutation of α1G T-type calcium channels in mice does not change anesthetic requirements for loss of the righting reflex and minimum alveolar concentration but delays the onset of anesthetic induction. Anesthesiology 106, 1177–1185 (2007).

25. Talley, E. M. et al. Differential Distribution of Three Members of a Gene Family Encoding Low Voltage-Activated (T-Type) Calcium Channels. J. Neurosci. 19, 1895–1911 (1999).

26. Schiff, N. D., Nauvel, T. & Victor, J. D. Large-scale brain dynamics in disorders of consciousness. Current Opinion in Neurobiology 25, 7–14 (2014).

27. Song, I. et al. Role of the α1G T-Type Calcium Channel in Spontaneous Absence Seizures in Mutant Mice. J. Neurosci. 24, 5249–5257 (2004).

28. Lee, J. et al. Sleep spindles are generated in the absence of T-type calcium channel-mediated low-threshold burst firing of thalamocortical neurons. Proceedings of the National Academy of Sciences 110, 20266–20271 (2013).

29. McCORMICK, D. A. & Pape, H.-C. Properties of a hyperpolarization-activated cation current and its role in rhythmic oscillation in thalamic relay neurones. The Journal of physiology 431, 291–318 (1990).

30. Saalmann, Y. B. Intralaminar and medial thalamic influence on cortical synchrony, information transmission and cognition. Front Syst Neurosci 8, (2014).

31. Giber, K. et al. A subcortical inhibitory signal for behavioral arrest in the thalamus. Nat Neurosci 18, 562–568 (2015).

32. Gummadavelli, A. et al. Thalamic stimulation to improve level of consciousness after seizures: Evaluation of electrophysiology and behavior. Epilepsia 56, 114–124 (2015).

33. He, J. h., et al. Decreased functional connectivity between the mediodorsal thalamus and default mode network in patients with disorders of consciousness. Acta Neurol Scand n/a-n/a (2014) doi:10.1111/ane.12299.

34. Choi, S., Yu, E., Lee, S. & Llinás, R. R. Altered thalamocortical rhythmicity and connectivity in mice lacking CaV3.1 T-type Ca2+ channels in unconsciousness. PNAS 201420983 (2015) doi:10.1073/pnas.1420983112.

35. Bastos, A. M. et al. Neural effects of propofol-induced unconsciousness and its reversal using thalamic stimulation. eLife 10, e60824 (2021).

36. Hwang, E., Kim, S., Shin, H.-S. & Choi, J. H. The forced walking test: A novel test for pinpointing the anesthetic-induced transition in consciousness in mouse. Journal of Neuroscience Methods 188, 14–23 (2010).

37. Cheong, E. et al. Deletion of phospholipase C β4 in thalamocortical relay nucleus leads to absence seizures. PNAS 106, 21912–21917 (2009).

38. Poulet, J. F., Fernandez, L. M., Crochet, S. & Petersen, C. C. Thalamic control of cortical states. Nature neuroscience 15, 370–372 (2012).

39. Schiff, M. L. & Reyes, A. D. Characterization of thalamocortical responses of regular-spiking and fast-spiking neurons of the mouse auditory cortex in vitro and in silico. Journal of Neurophysiology 107, 1476–1488 (2012).

40. Destexhe, A. Self-sustained asynchronous irregular states and Up–Down states in thalamic, cortical and thalamocortical networks of nonlinear integrate-and-fire neurons. J Comput Neurosci 27, 493–506 (2009).

41. Llinás, R. R. & Steriade, M. Bursting of Thalamic Neurons and States of Vigilance. Journal of Neurophysiology 95, 3297–3308 (2006).

42. McInnes, L., Healy, J. & Melville, J. Umap: Uniform manifold approximation and projection for dimension reduction. *arXiv preprint arXiv:1802.03426* (2018).

43. Gangadharan, G. et al. Medial septal GABAergic projection neurons promote object exploration behavior and type 2 theta rhythm. Proceedings of the National Academy of Sciences 113, 6550–6555 (2016).

44. Lee, J.-H. et al. The rostroventral part of the thalamic reticular nucleus modulates fear extinction. Nat Commun 10, 4637 (2019).

45. Lee, S. et al. Bidirectional modulation of fear extinction by mediodorsal thalamic firing in mice. Nat Neurosci 15, 308–314 (2012).

46. Courtiol, E. & Wilson, D. A. Thalamic olfaction: characterizing odor processing in the mediodorsal thalamus of the rat. Journal of Neurophysiology 111, 1274–1285 (2014).

47. Paydar, A. et al. Extrasynaptic GABAA receptors in mediodorsal thalamic nucleus modulate fear extinction learning. Molecular Brain 7, 39 (2014).

48. Colavito, V., Tesoriero, C., Wirtu, A. T., Grassi-Zucconi, G. & Bentivoglio, M. Limbic thalamus and state-dependent behavior: The paraventricular nucleus of the thalamic midline as a node in circadian timing and sleep/wake-regulatory networks. Neuroscience & Biobehavioral Reviews (2014) doi:10.1016/j.neubiorev.2014.11.021.

49. Sarter, M. & Bruno, J. P. Cortical cholinergic inputs mediating arousal, attentional processing and dreaming: differential afferent regulation of the basal forebrain by telencephalic and brainstem afferents. Neuroscience 95, 933–952 (1999).

50. Cassidy, R. M. & Gale, K. Mediodorsal thalamus plays a critical role in the development of limbic motor seizures. The Journal of neuroscience 18, 9002–9009 (1998).

51. Deleuze, C. et al. T-Type Calcium Channels Consolidate Tonic Action Potential Output of Thalamic Neurons to Neocortex. J. Neurosci. 32, 12228–12236 (2012).

52. Tscherter, A. et al. Minimal alterations in T-type calcium channel gating markedly modify physiological firing dynamics. The Journal of Physiology 589, 1707–1724 (2011).

53. Mease, R. A., Krieger, P. & Groh, A. Cortical control of adaptation and sensory relay mode in the thalamus. PNAS 111, 6798–6803 (2014).

54. Hildebrand, M. E. et al. Functional Coupling between mGluR1 and Cav3.1 T-Type Calcium Channels Contributes to Parallel Fiber-Induced Fast Calcium Signaling within Purkinje Cell Dendritic Spines. J. Neurosci. 29, 9668–9682 (2009).

55. Crunelli, V., David, F., Leresche, N. & Lambert, R. C. Role for T-type Ca2+ channels in sleep waves. Pflügers Archiv-European Journal of Physiology 466, 735–745 (2014).

56. Shao, J. et al. Cav3.1-driven bursting firing in ventromedial hypothalamic neurons exerts dual control of anxiety-like behavior and energy expenditure. Mol Psychiatry 27, 2901–2913 (2022).

57. Guido, W., Lu, S. M. & Sherman, S. M. Relative contributions of burst and tonic responses to the receptive field properties of lateral geniculate neurons in the cat. J. Neurophysiol. 68, 2199–2211 (1992).

58. Montemurro, M. A. et al. Role of Precise Spike Timing in Coding of Dynamic Vibrissa Stimuli in Somatosensory Thalamus. Journal of Neurophysiology 98, 1871–1882 (2007).

59. Bereshpolova, Y. et al. Getting drowsy? Alert/nonalert transitions and visual thalamocortical network dynamics. The Journal of Neuroscience 31, 17480–17487 (2011).

60. Paz, J. T. et al. Closed-loop optogenetic control of thalamus as a tool for interrupting seizures after cortical injury. Nat Neurosci 16, 64–70 (2013).

61. Abdelaal, M. S. et al. Dysfunction of parvalbumin-expressing cells in the thalamic reticular nucleus induces cortical spike-and-wave discharges and an unconscious state. Brain Communications 4, fcac010 (2022).

62. Lory, P., Nicole, S. & Monteil, A. Neuronal Cav3 channelopathies: recent progress and perspectives. Pflugers Arch 472, 831–844 (2020).

63. Ehlers, C. L. & Slawecki, C. J. Effects of chronic ethanol exposure on sleep in rats. Alcohol 20, 173–179 (2000).

64. Carden, W. B. et al. Chronic ethanol drinking reduces native T-type calcium current in the thalamus of nonhuman primates. Brain Research 1089, 92–100 (2006).

65. Clasadonte, J., McIver, S. R., Schmitt, L. I., Halassa, M. M. & Haydon, P. G. Chronic Sleep Restriction Disrupts Sleep Homeostasis and Behavioral Sensitivity to Alcohol by Reducing the Extracellular Accumulation of Adenosine. J. Neurosci. 34, 1879–1891 (2014).

66. Naassila, M., Ledent, C. & Daoust, M. Low Ethanol Sensitivity and Increased Ethanol Consumption in Mice Lacking Adenosine A2A Receptors. J. Neurosci. 22, 10487–10493 (2002).

67. Allan, A. M. & Harris, R. A. Acute and chronic ethanol treatments alter GABA receptor-operated chloride channels. Pharmacology Biochemistry and Behavior 27, 665–670 (1987).

68. Suryanarayanan, A. et al. Subunit compensation and plasticity of synaptic GABAA receptors induced by ethanol in α4 subunit knockout mice. Frontiers in neuroscience 5, (2011).

69. Koob, G. F. et al. Neurocircuitry Targets in Ethanol Reward and Dependence. Alcoholism: Clinical and Experimental Research 22, 3–9 (1998).

70. Schulteis, G. & Liu, J. Brain reward deficits accompany withdrawal (hangover) from acute ethanol in rats. Alcohol 39, 21–28 (2006).

71. Feng, L., Kwon, O., Lee, B., Oh, W. C. & Kim, J. Using mammalian GFP reconstitution across synaptic partners (mGRASP) to map synaptic connectivity in the mouse brain. Nat. Protocols 9, 2425–2437 (2014).

72. Kohtoh, S. et al. Algorithm for sleep scoring in experimental animals based on fast Fourier transform power spectrum analysis of the electroencephalogram. Sleep and Biological Rhythms 6, 163–171 (2008).

73. Stephenson, R., Caron, A. M., Cassel, D. B. & Kostela, J. C. Automated analysis of sleep–wake state in rats. Journal of neuroscience methods 184, 263–274 (2009).

