## supplementary Data for "Mediodorsal thalamic nucleus mediates resistance to ethanol through Cav3.1 T-type Ca^2+^ regulation of neural activity"

$Ca_v3.1$ T-type calcium channel, mediodorsal thalamic nucleus, ethanol resistance, loss of consciousness, tonic and burst firing, optogenetic/electrical stimulation

**CONTENT**

Supplementary figures: 12
Supplementary method
Supplementary references

#### Supplementary Figures

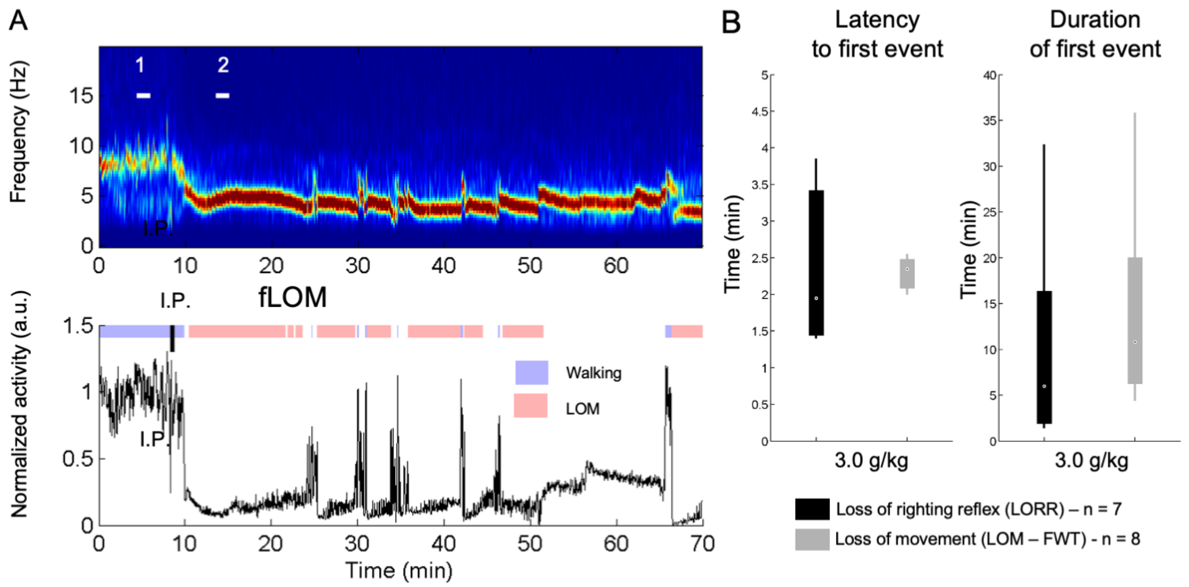


**Fig. S1: The forced walking task (FWT) is an uninterrupted assessment of the mouse sedative state equivalent to the loss of righting reflex.**

(A) Representative recording for the forced walking task (FWT) with a 10 min baseline walking and 60 min observation after I.P. injection; EEG power spectrogram depicting the typical theta rhythm (4-6Hz) associated with the ethanol hypnotic state(upper panel), normalized root mean square of accelerometer activity (RMS; bottom panel) and interpreted mouse activity using the normalized RMS (see experimental procedures for details) highlighted in blue (Walking) and red (loss of motion; LOM) on a constantly running treadmill (6 cm/s) with plexiglass walls and no resting area; the black tick indicates the time of intraperitoneal (I.P.) injection of ethanol (3.0 g/Kg);

(B) Latency to and duration of an episode of LORR or LOM shows less variability in FWT assessment with similar values indicating equivalence in assessment power.

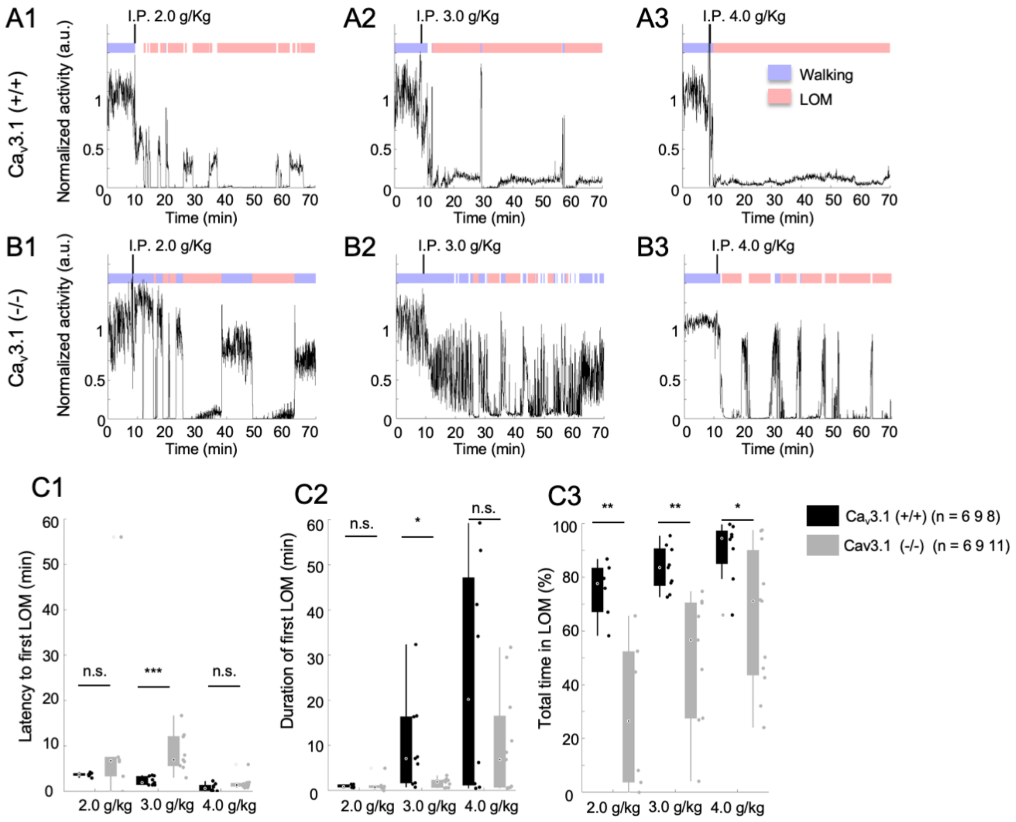


**Fig. S2: Ca$_v$3.1 mutant mice show a dose-dependent ethanol resistance.**

(A) Representative motor activity and loss of movement (LOM) over time for Ca$_v$3.1(+/+) mice post I.P. injection of 2.0, 3.0 and 4.0 g/Kg.

(B) Representative motor activity and loss of movement (LOM) over time for Ca$_v$3.1 (-/-) mice post I.P. injection of 2.0, 3.0 and 4.0 g/Kg.

(C) Latency to first LOM (fLOM; C1), duration of the first LOM (C2) and total time spent in LOM state (C3) over a recording duration of 1hour following 2.0, 3.0 and 4.0 g/Kg I.P injection of ethanol in Ca$_v$3.1 (+/+) and Ca$_v$3.1 (-/-) mice; data is represented as boxplot. Group and dose effects were assessed using a two-way ANOVA. Post-hoc group comparison was performed using a Holm-Sidak correction for multiple comparison. * is for $p<0.05$, ** is for $p<0.01$, *** is for $p<0.001$ and n.s. is for non-significant.

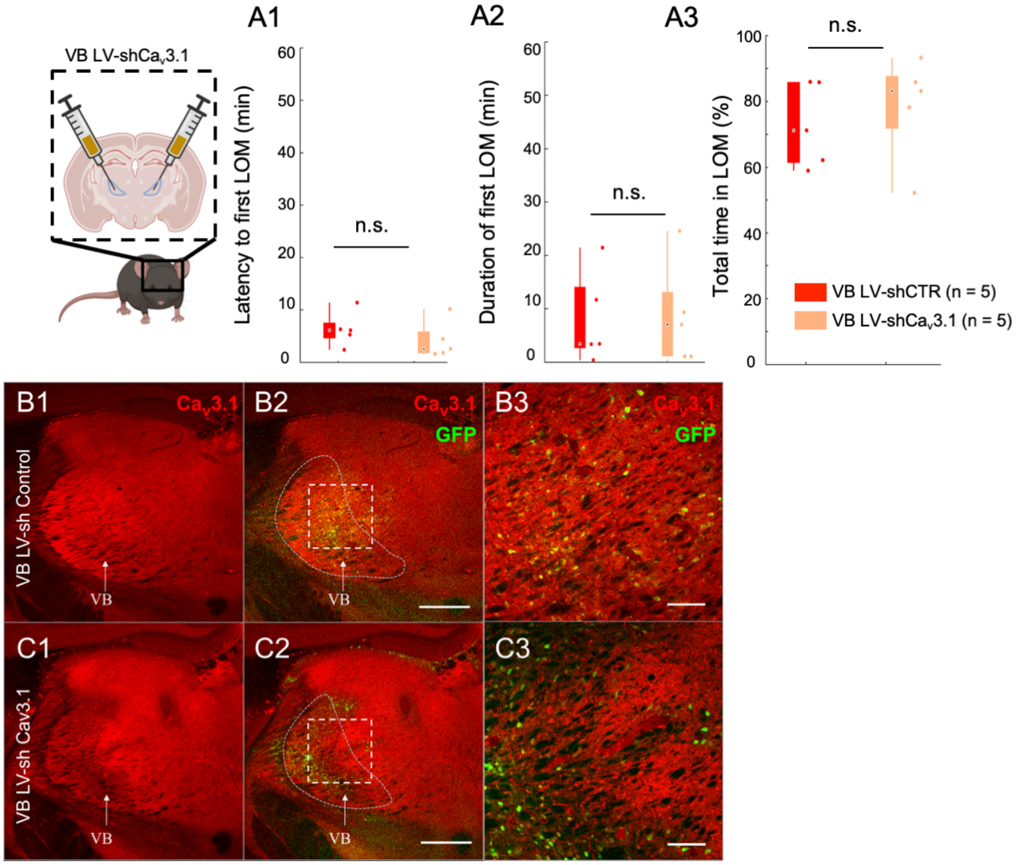


**Fig. S3: Mice with ventrobasal nucleus Ca$_v$3.1 knock-down did not show increased ethanol resistance.**

(A) Latency to first LOM (fLOM; A1), duration of the first LOM (A2) and total time spent in LOM state (A3) over a recording duration of 60 min post I.P. injection of 3.0g/Kg I.P. injection of EtOH in lentivirus-shControl and shCa$_v$3.1 knock-down mice for MD ; data is represented as boxplot. *, ** and n.s are for $p<0.05$, $p<0.01$ and non-significant, respectively.

(B) Representative brain coronal section stained for Ca$_v$3.1 anti-body (B1) showing the wide thalamic expression of the t-type calcium channel in the VB lentivirus(LV)-shControl injected mice; Ca$_v$3.1 and GFP merging (B2) and higher magnification of the white dashed square in B1-B2 (B3). Scale bars in (B2) and (B3) indicate 500 µm and 100 µm, respectively.

(C) Representative brain coronal section stained for Ca$_v$3.1 anti-body (C1) showing the reduction in t-type calcium channel expression in the VB LV-shCa$_v$3.1 injected mice; Ca$_v$3.1 and GFP merging (C2) and higher magnification of the white dashed square in C1-C2 (C3). Scale bars in (C2) and (C3) indicate 500 µm and 100 µm, respectively.

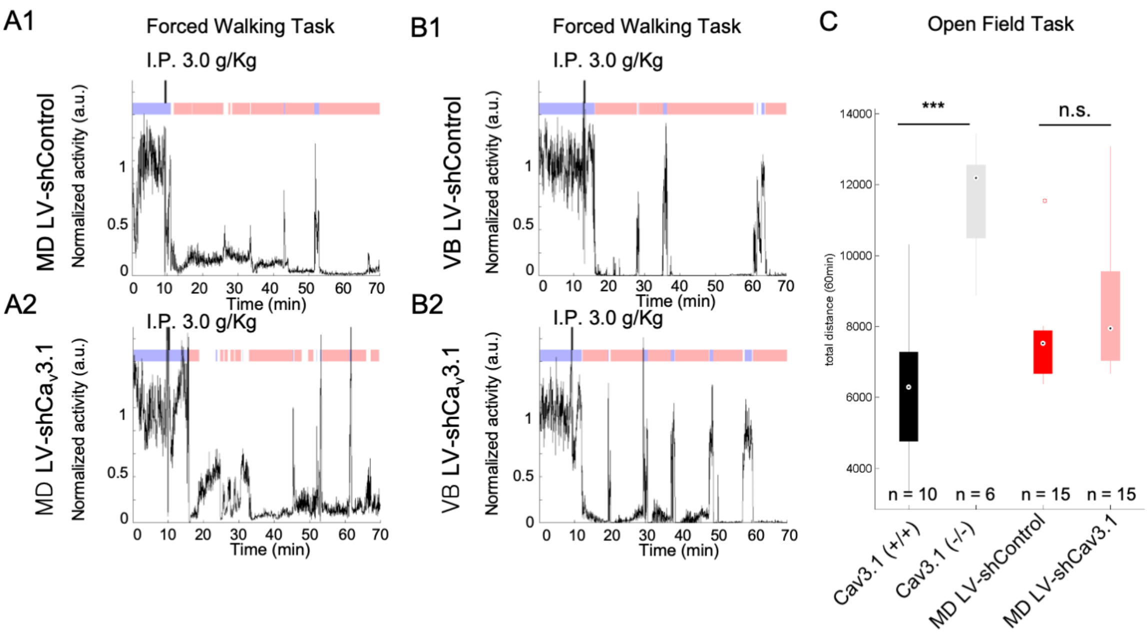


**Fig. S4: Representative activity of mice with Ca$_v$3.1 knock-down in MD and VB.**

(A) Representative activity separated in walking (blue) and LOM (red) over time for MD Lentivirus-shControl (A1) and MD Lentivirus-shCa$_v$3.1(A2).

(B) Representative activity separated in walking (blue) and LOM (red) over time for VB Lentivirus-shControl (B1) and VB Lentivirus-shCa$_v$3.1(B2).

(C) Boxplot of total distance moved over 30 min for Ca$_v$3.1(+/+), Ca$_v$3.1(-/-), MD LV-shControl and MD LV-shCa$_v$3.1 mice; * is for $p<0.05$, ** is for $p<0.01$ and *** is for $p<0.001$.

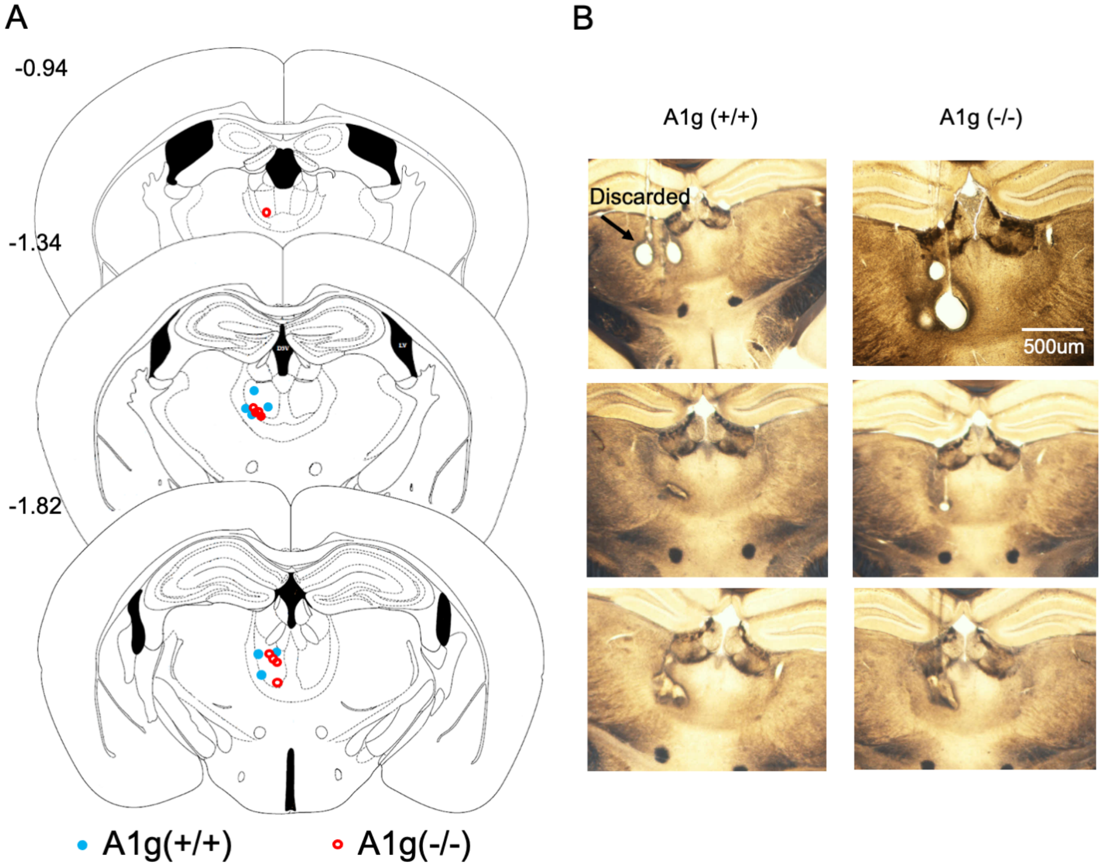


**Fig. S5: Representative positioning of single unit recording in the mouse MD.**

(A) Summary localization of tetrode ending tips in the MD using post-mortem histological analysis. Each tetrode wire applied a 10sec 1mA current to create an electro lesion at the end tip prior euthanasia.

(B) Representative electrolesion in the central (B1) and ventral (B2) MD.

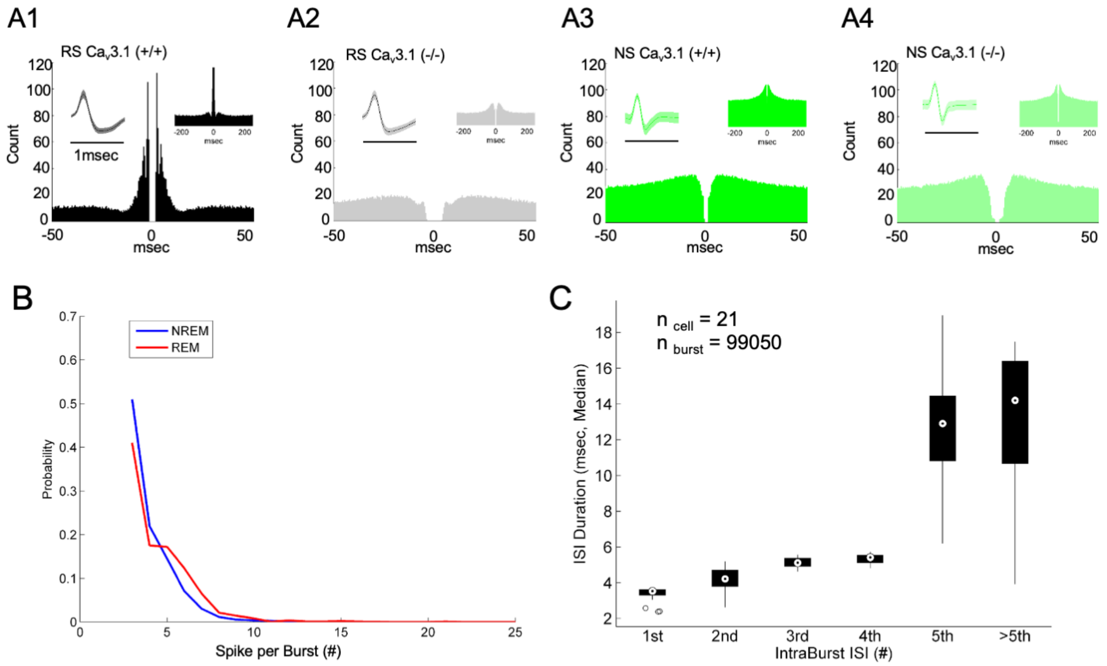


**Fig. S6: Representative putative neurons and burst firing properties in MD**

(A) Representative waveform and auto-cross correlogram of a regular spiking (wider spike waveform), expected to represent putative excitatory neurons from $Ca_v$3.1 (+/+) (A1) and $Ca_v$3.1 (-/-) (A2). Representative waveform and auto-cross correlogram of a narrow spiking neuron, expected to represent putative inhibitory neurons (central thalamus or projecting parvalbumin neurons from the thalamic reticular nucleus) from $Ca_v$3.1 (+/+) (A3) and $Ca_v$3.1 (-/-) (A4).

(B) Probability of spike per burst estimated during non-rapid eye movement (NREM) and rapid eye movement (REM) sleep in $Ca_v$3.1 WT mice.

(C) Boxplot of intra-burst inter-spike interval for the five first observed spikes and above. On average the 4 first spikes ISI remains below 6ms (intra-burst frequency >166 Hz).

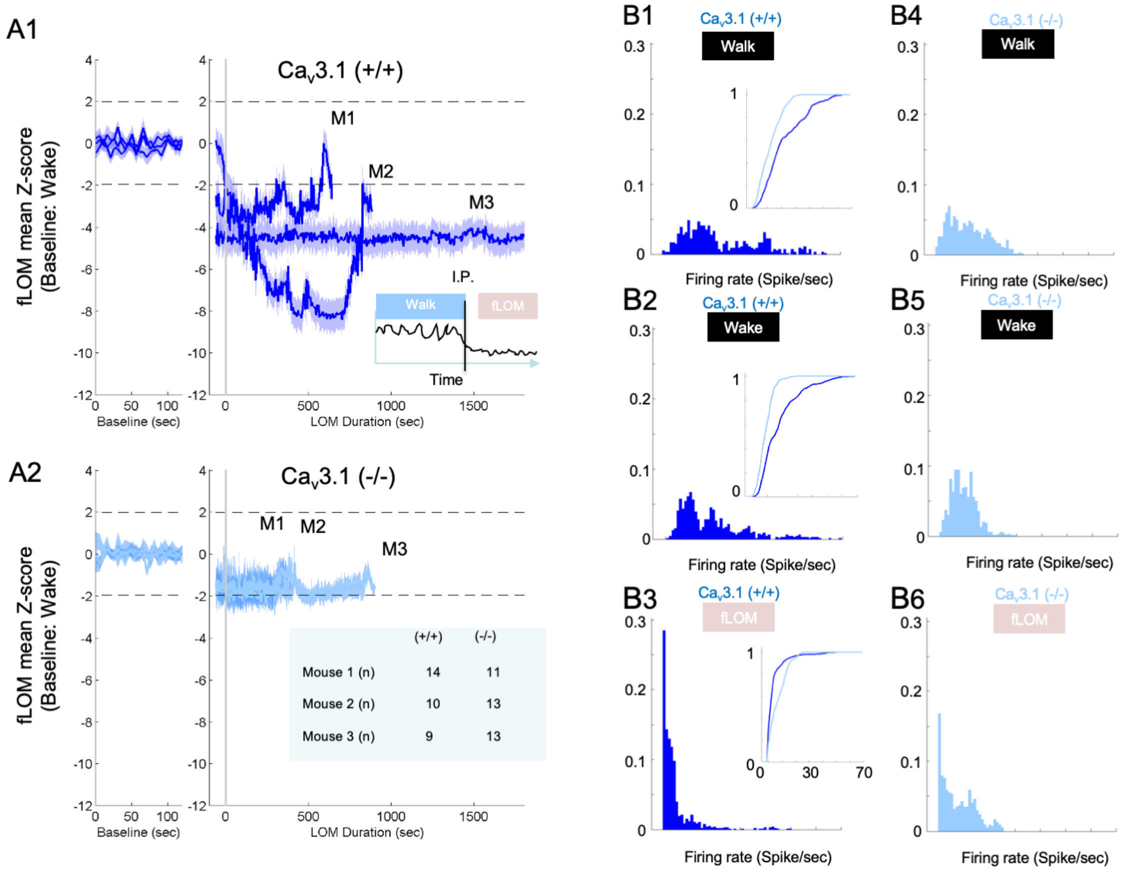


**Fig. S7: MD neurons activity remains within wakefulness level during fLOM in $Ca_v3.1$ mutant.**

(A) Mean MD firing Z-score traces (normalized with respect to wakeful state) during walking and fLOM for WT (A1) and mutant mice (A2); data is represented as mean ± s.e.m.; dash lines represent Z-score significance level Z = +/- 1.96.

(B) Distribution of tonic firing rate of MD RS neurons of $Ca_v3.1$(+/+) during walk (B1), wake (B2) and fLOM (B3) and $Ca_v3.1$(-/-) during walk (B4), wake (B5) and fLOM (B6) mice; inner panels in B1, B2 and B3 represents the comparative plot of cumulative distribution of tonic firing between wild type and mutant MD RS neurons.

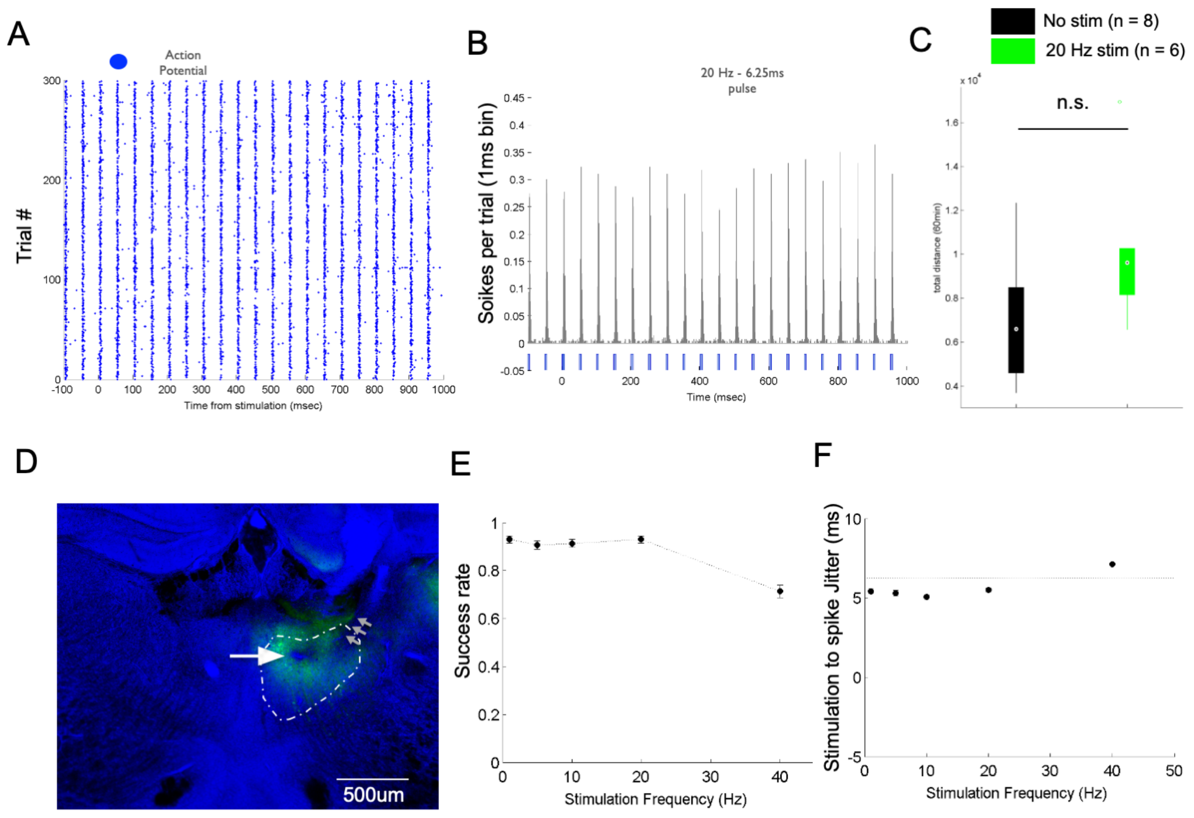


**Fig. S8: Optogenetic stimulation-induced sustained response in MD**

(A) Evoked spike response during optogenetic stimulation of MD RS neutron for 300 trial/stimulation at 20Hz (pulse width = 6.25 sec);

(B) Peri-stimulus histogram (PSTH) of the evoked response shown in using a 1 msec-bin.

(C) Boxplot of total distance moved over 30 min for no stimulation and tonic-like stimulation; * is for $p<0.05$, ** is for $p<0.01$ and *** is for $p<0.001$.

(D) Coronal section showing lesion at the site of tetrode recording (white arrow) and puncture due to optical fiber cannula (grey arrows); green fluorescence indicates GFP expression after syn-AAV9-ChR2-sfGFP virus infection.

(E) Success rate of stimulation estimated for protocol of different frequencies stimulation; data is represented as mean ± s.e.m.

(F) Estimation of delay between laser pulse onset (time equal 0) and the first evoked spikes for different frequency stimulation; data is represented as mean ± s.e.m.; the gray horizontal line indicates the end of the laser pulse.

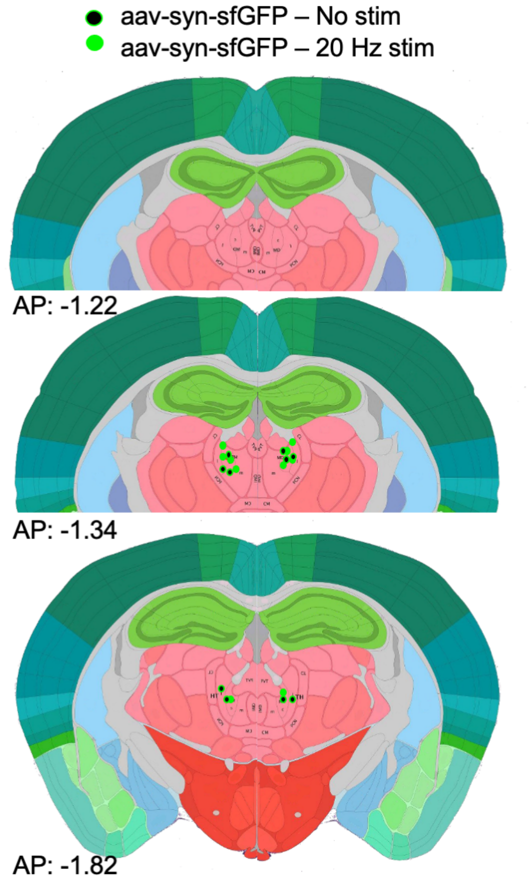


**Fig. S9: Positioning summary of optic fiber cannula used for optogenetic stimulation.**

(A) Summary of optic fiber cannula positioning used for bilateral optogenetic stimulation of $Ca_v3.1$(+/+) mice injected with syn-AAV9-ChR2-sfGFP.

(B) Representative GFP images of ChR2-sfGFP expression and optic fiber cannula position (white arrows).

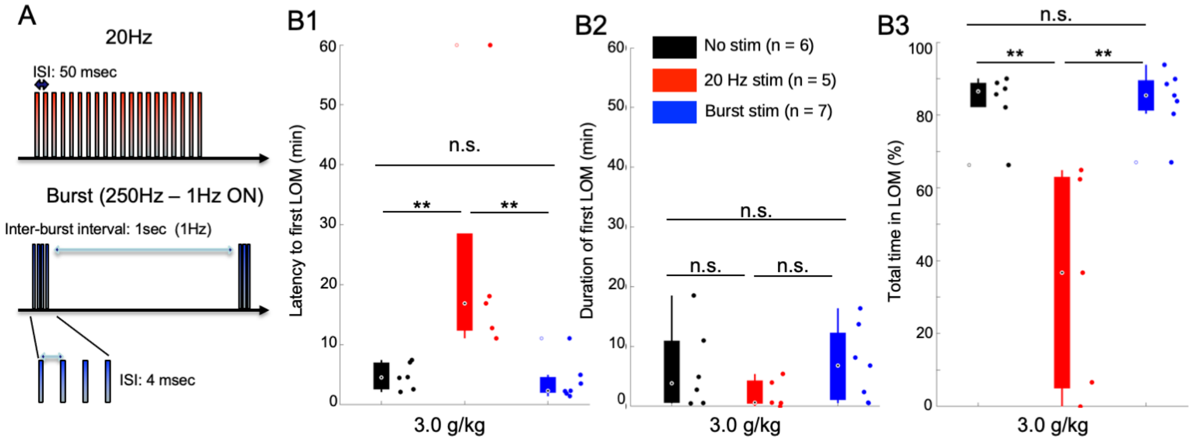


**Fig. S10: Electrical 20 Hz stimulation of MD increases ethanol resistance in WT mice.**

(A) Electric stimulation protocol for the 20Hz (tonic-like) and burst-like stimulation.

(B) Latency to first LOM (B1), duration of the first LOM (B2) and total time spent in LOM state (B3) over a recording duration of one hour post I.P. injection of 3.0g/Kg of ethanol for optogenetic stimulation. * is for $p<0.05$, ** is for $p<0.01$, *** is for $p<0.001$ and n.s. is for non-significant.

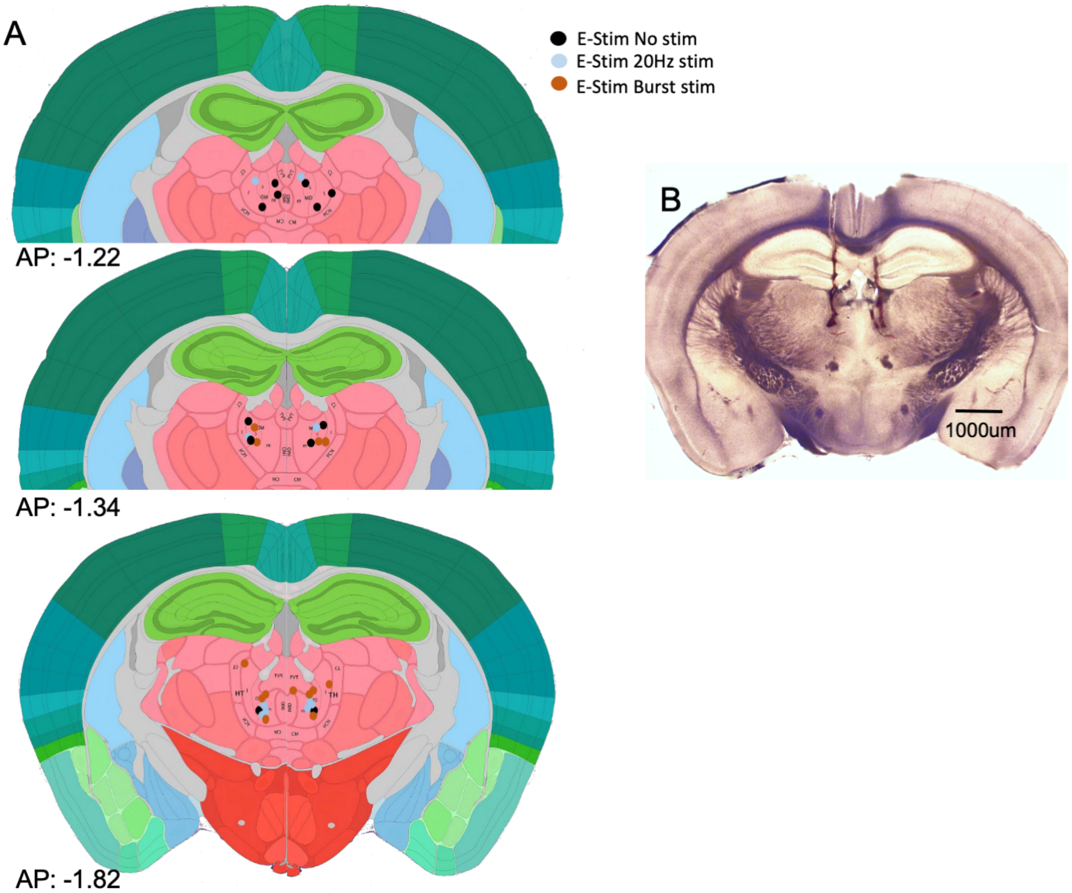


**Fig. S11: Positioning summary of electrodes used for electric stimulation.**

(A) Summary of positioning used for bilateral bipolar electrical stimulation of $Ca_v3.1$(+/+) mice.

(B) Representative bright field images of stimulation electrode positioning.

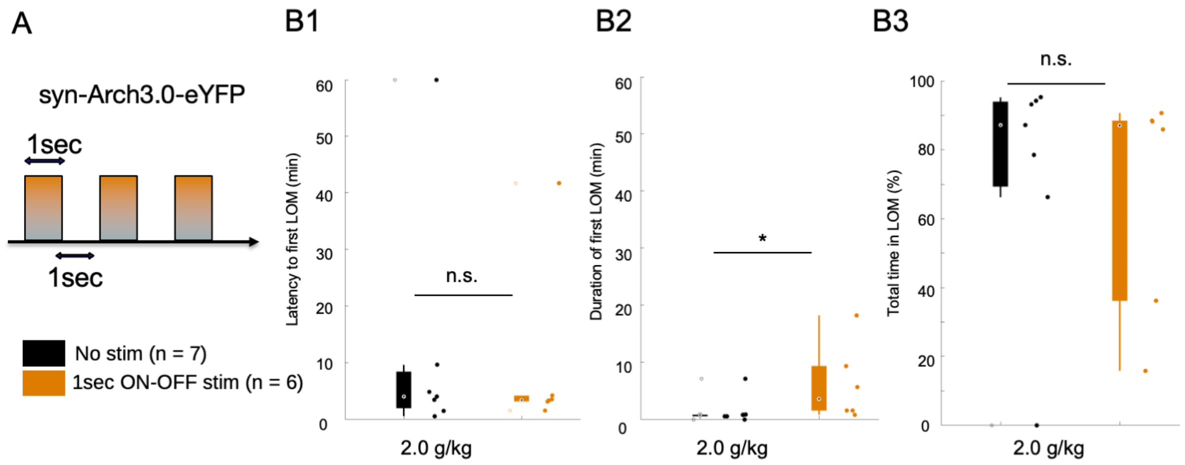


**Fig. S12: 1sec ON-OFF optogenetic inhibition of MD increases LOM duration in WT mice.**

(A) WT mice were transduced with an AAV-syn-ARCH3.0-eYFP to express archaerhodopsin and implanted with bilateral optic fibers as in the channelrhodopsin experiment. A phasic stimulation of 1secon ON-OFF was then performed to promote inhibition and possible rebound burst [60,61]. A lower dose of ethanol (2.0 g/Kg) was used to test the prolongation of the hypnotic effect by the phasic inhibition of MD.

(B) Latency to (B1), duration of (B2) and total time in LOM

### Supplementary Methods

**Surgery for electrophysiological recordings**. The surgical implantation of electrodes (EEG, EMG and/or tetrode Microdrive) and virus injection procedures were performed under 0.2% tribromoethanol (Avertin) anesthesia (20 mL/kg i.p.). Following anesthetic administration, mice (11-week-old for electrode implantation; 10 weeks for virus injection) were fixed in a stereotaxic device (David Kopf Instruments). For chronic recording of EEG and EMG, a stainless-steel screw electrode was fixed into the skull over the right parietal hemisphere and an uncoated stainless-steel wire was tied to the nuchal muscle, respectively. For in vivo freely moving single unit recording, we used a Harlan 4 Drive (Neuralynx inc.) mounted with 3 to 4 tetrode wires inserted to the caudal region of the right mediodorsal thalamic nucleus (anteroposterior, −1.4; lateral, +0.4; depth: 3.2 mm). Single tetrode wires were prepared from 4 twisted nichrome-formvar/PAC wires (Kanthal precision technology, OD 0.0127 mm) and gold plated to achieve an impedance range of 150-400kΩ (1kHz, in saline solution). A period of  7 days was given to allow a complete recovery from the surgical procedure.

**EEG/EMG recordings**. EEG signals were amplified and band-pass filtered in the range 0.1-100 Hz. EMG signals were high-pass filtered at 70 Hz. All recordings were digitized at a sampling rate of 1kHz (Grass Amplifiers, pClamp 9.2-Molecular devices) or at 32kHz (Cheetah 6.5-Neuralynx) and downsampled in post-processing.

**Single Unit recording, sorting and analysis**. Electrophysiological data obtained from tetrodes bundles were acquired using Digital Lynx hardware and Cheetah 6.5 (Neuralynx) at a sampling frequency of 32 kHz. Online band-pass filtering (LFP for spike sorting: 600-6000 Hz; EEG: 0.5-70 Hz; EMG: 70-4000 Hz) and spike sorting was performed using cheetah 6.5. Off-line spike clustering and sorting was performed semi automatically using KlustaKwik (K.D. Harris, http://klustakwik.sourceforge.net) and MClust 3.5 (A.D. Redish, http://redishlab.neuroscience.umn.edu) in Matlab (R) (the MathWorks, inc.) or SpikeSort3D 2.5. The time stamps or spike trains associated with each identified single unit were analyzed using a customized algorithm through Matlab (R). Single unit characterization was performed by means of using inter-spike interval distribution (ISI), cross- and autocross-correlation histograms (e.g. bursting index, bursting mode, spectral distribution) , inter- and intra-burst property analysis (e.g. intra-burst ISI, number of spikes per burst, burst spike rate) and associated spike waveform indices (e.g. peak, peak-to-valley spike width, first and second principal component). Population spiking was analyzed by means of peri-event histogram, normalized cross-correlation pairs and phase coherency, using chronux toolbox (chronux.org) and custom-made codes.

The bursting index was derived as described in (Royer et al. 2012). Namely, the burst index was estimated from the spike auto-correlogram (1-ms bin size) by subtracting the mean value between 40 and 50 ms (baseline) from the peak measured between 0 and 10 ms. Positive burst amplitudes were normalized to the peak and negative amplitudes were normalized to the baseline to obtain indexes ranging from −1 to 1.

**Sleep Monitoring and Staging**. Sleep scoring was based on the EEG and EMG recordings obtained from a period of 6 hours recorded in the second phase of the light cycle (12:00-18:00). We used a custom-made automatic sleep scoring system based on two previously described scoring methods for rodents [72,73] and organized a voting scheme for the final staging decision. All sleep scores were visually inspected and corrected by a sleep specialist.

**Immunohistochemistry.** Sections of perfused mouse brain (5% formaldehyde) were intensively washed with phosphate buffer (0.1 M) and then treated with a blocking solution containing 3% normal donkey serum (Millipore) and 0.2% Triton-X (Sigma) for 40 min at room temperature. The following primary antibodies diluted in phosphate buffer were used: anti-$Ca_v$3.1 antibody (rabbit, 1:200; Alomone Labs), anti-NeuN antibody (mouse, 1:500; millipore) and anti-calbindin D-28k antibody (mouse, 1:3000; Swant). After primary antibody incubation (1 day at room temperature), sections were treated with secondary antibodies labeled with fluorescent dye (Cy3 or Cy5; 1:500, 2h at room temperature; Jackson). Sections with fluorescent staining were mounted in a mounting solution (VECTASHIELD with DAPI; Vector Laboratories, H-1200). Photographs were taken using either a microscope (Nikon Eclipse-Ti) or a FluoView FV1000 confocal laser scanning system (Olympus). When necessary, brightness and contrast were adjusted using the FluoView client program applied to whole images only.

**Uniform Manifold Approximation and Projection (UMAP)**

In order to provide a visual representation of the various brain states recorded in $Ca_v$3.1 wild type and mutant mice, we combined the tonic firing rate, burst firing rate and burst event rate into a reduced manifold representation using the UMAP method[42]. The version of MATLAB implementation was used with a fixed seed input.

**Archaerhodopsin-mediated inhibition of MD neurons**

For our phasic inhibition experiment, 16 mice were injected in MD with aav5.hSyn.eArch3.0-eYFP (University of North Carolina, Vector Core). 3 mice were discarded post histological analysis due to low viral expression. This construct was favored over the halorhodopsin channel due to the long duration of the stimulation intended (60 min, 1sec pulse with a duty cycle of 50%, 0r 1 s ON-OFF sequence) and low toxicity. The mice were implanted with optic fiber guides (125 μm core diameter, Doric Lenses inc.) positioned at 30 degree angle from the transverse plane. The mice were given 2~3 weeks to recover and to allow for the viral expression. For Arch-mediated inhibition we used a 532 nm (Green, MGL-S-532-OEM, Changchun New Industries Optoelectronics Technology Co., Ltd) laser to deliver at ~2mW to each fiber guide through a patch cord (SMA end-to-end; Thorlabs Inc.). These mice were then randomly assigned to a no stimulation (n = 7) and a 1 s ON-OFF stimulation group (n = 6). The mice received the stimulation immediately after being placed in the treadmill, then received the i.p. injection of 3.0 g/kg of ethanol as in other experiments.

#### Supplemental References

Kohtoh, S., Taguchi, Y., Matsumoto, N., Wada, M., Huang, Z.-L., and Urade, Y. (2008). Algorithm for sleep scoring in experimental animals based on fast Fourier transform power spectrum analysis of the electroencephalogram. Sleep Biol. Rhythms *6*, 163–171.

Stephenson, R., Caron, A.M., Cassel, D.B., and Kostela, J.C. (2009). Automated analysis of sleep–wake state in rats. J. Neurosci. Methods *184*, 263–274.

Royer, S., Zemelman, B. V., Losonczy, A., Kim, J., Chance, F., Magee, J. C., & Buzsáki, G. (2012). Control of timing, rate and bursts of hippocampal place cells by dendritic and somatic inhibition. *Nature neuroscience*, *15*(5), 769-775.
